# Cytoskeletal-nuclear control of alveolar fibroblast identity directs lung regeneration

**DOI:** 10.64898/2026.08.17.745350

**Authors:** Dakota L. Jones, Sarah E. Schaefer, Michael P. Morley, Kazushige Shiraishi, Parisha Shah, Ricardo A. Linares-Saldana, Yun Ying, Ullas V. Chembazhi, Su Zhou, Rajan Jain, Edward E. Morrisey

## Abstract

Respiratory mechanics direct cell fate in the lung, but the mechanisms by which these mechanical signals are sensed and transmitted to the nucleus to control cell state remain unclear. We paired in vivo perturbations of respiratory mechanics with single-cell genomics and found that alveolar fibroblasts are highly sensitive to physical changes in their microenvironment, exhibiting persistent shifts in their transcriptional identity after injury. Surprisingly, transmission of these signals through the nuclear envelope was not essential for maintaining transcriptional or epigenetic stability during homeostasis. However, severing mechanical-nuclear signaling promoted the normalization of alveolar fibroblast identity after acute injury, resulting in improved epithelial regeneration and reduced dysplastic remodeling. These studies reveal the importance of mechanical-nuclear signaling in the regulation of alveolar cell identity and function and reveal that targeting this complex can enhance tissue regeneration.

## Introduction

Efficient lung regeneration requires coordinated communication between epithelial and mesenchymal cells within the alveolus. Alveolar fibroblasts play a central role in this process by maintaining a supportive niche that sustains alveolar type II (AT2) epithelial progenitor cell function through paracrine signals such as Wnt and Fgf^1–4^. These niche-supportive functions depend on a stable alveolar fibroblast cell identity defined by a transcriptional and functional program that preserves alveolar structure and regenerative capacity^5,6^. Following injury, alveolar fibroblasts transiently activate to support repair. However, when this activation persists or is dysregulated, alveolar fibroblasts lose their niche identity, adopt pathogenic phenotypes, and contribute to aberrant remodeling characteristic of chronic lung disease^5–7^. While the pathways regulating fibroblast activation have been extensively studied, how alveolar fibroblast identity is maintained or destabilized within the alveolar microenvironment remains poorly understood.

The lung alveolus is a mechanically active structure that expands and contracts with each breath. Mechanical forces arising from respiratory mediated stretch, altered matrix stiffness, and disrupted tissue architecture are increasingly recognized as key regulators of alveolar cell behavior and drivers of chronic lung pathology and disease^8–12^. Previous work has shown that lung fibroblasts cultured on stiff substrates that model fibrotic lung tissue spontaneously lose their identity, which is accompanied by alterations in nuclear morphology and chromatin organization that promote persistent, disease-like transcriptional states^13,14^. These observations suggest that mechanical signaling transmitted to the nucleus induces changes in chromatin structure and transcriptional programs, thereby destabilizing fibroblast identity and impairing their regenerative function. However, the mechanisms by which nuclear mechanotransduction governs fibroblast cell identity in vivo remain unresolved.

Cells transmit cytoskeletal forces to the nucleus through the LINC (Linker of Nucleoskeleton and Cytoskeleton) complex, composed of outer nuclear membrane Nesprins and inner nuclear membrane Sun1/2 proteins^15^. The LINC complex mechanically couples cytoskeletal tension to the nuclear lamina and the underlying chromatin, influencing lamin-associated domains (LADs), chromatin accessibility, heterochromatin formation, and gene expression^16^. Although canonical mechano-transducers such as Yap and Taz are known to regulate lung cell behavior^17–24^, their extensive crosstalk with Wnt, Notch, and TGF-beta, and other key pathways complicates interpretation of their mechanotransduction outputs^25^. Despite its importance in mechanical signaling in cells, the in vivo role of LINC-mediated mechanotransduction in tissue homeostasis and regeneration has remained largely unexplored.

To distinguish alveolar fibroblast responses from physiologic regeneration to maladaptive injury, we examined alveolar fibroblast responses in multiple models of lung injury and changes to respiratory mechanics. Single-cell transcriptomic analyses revealed that alveolar fibroblasts exhibit a persistent reduction in identity following bleomycin-induced lung injury, while simultaneously exhibiting a marked increase in mechanotransduction-related pathways. To test whether mechanotransduction and alveolar fibroblast identity are intrinsically linked, we used in vivo models to manipulate alveolar tissue mechanics, partial pneumonectomy (PNX) (gain-of-stretch) and bronchial ligation (loss-of-stretch). We found that disruption of alveolar tissue mechanics destabilized alveolar fibroblast identity, and that this destabilization is reversible upon recovery of homeostatic alveolar mechanics, indicating that injury-induced disruption of tissue mechanics can imprint a lasting, but potentially reversible, transcriptional state in alveolar fibroblasts. Disruption of cytoskeletal-nuclear force transmission by genetic deletion of Sun1 and Sun2, core proteins of the LINC complex, in alveolar fibroblasts restored their identity after bleomycin injury, re-establishing pro-regenerative Wnt/Fgf signaling, and improved epithelial regeneration. Together, these results identify nuclear mechanotransduction as a critical regulator of alveolar fibroblast identity and reveal that injury-induced mechanical forces can imprint a persistent memory that impairs tissue regenerative capacity.

## Results

### Persistent imbalance of alveolar fibroblast identity and mechanotransduction after injury

To investigate how acute lung injury affects the transcriptional state and identity of alveolar fibroblasts and other lung cell types, we used the intratracheal bleomycin model, which induces injury and transient fibrosis in the alveolar parenchyma that partially resolves over time (Fig. 1a, b). Following bleomycin administration, the stereotyped injury response includes diffuse alveolar damage and inflammation that peaks around day 14 and is considered largely resolved by day 28^26^. We re-analyzed our prior scRNA-seq dataset in which alveolar Pdgfra+ fibroblasts were tracked and transcriptionally profiled throughout injury and resolution. In parallel, we re-analyzed our companion scRNA-seq dataset generated from FACS-enriched CD45+ immune, CD31+ endothelial, and CD326+ epithelial cells collected from the same animals (Supp Fig. 1). After annotating all cell types in the merged dataset (Supp Fig. 1a), we performed differential expression analysis within each cell type at day 14 and day 28 relative to sham. Across the lung, alveolar fibroblasts exhibited the most robust transcriptional response, as measured by the total number of differentially expressed genes, with a striking number of differentially expressed genes persisting into the resolution phase at day 28 (Fig. 1d). Within the Pdgfra-lineage dataset, dimensionality reduction revealed distinct UMAP clusters driven primarily by time after bleomycin injury (Fig. 1e). Notably, at day 28, despite the tissue entering the resolution phase, we observed transcriptionally distinct subpopulations of Pdgfra+ alveolar fibroblasts that failed to return to their resting state, paralleling the persistence of dysplastic lung architecture at this time point (Fig. 1). Importantly, this persistence was not explained by sustained expansion of injury-induced fibroblast states, such as Cthrc1+ fibrotic fibroblasts (Cthrc1+ Spp1+ Postn+) or proliferating fibroblasts (Mki67+ Top2a+ Cdk1+), with only minor contributions from inflammatory fibroblasts (Saa3+ Mt2+ Lcn2+) (Supp Fig. 2a-c), suggesting that alveolar fibroblasts instead retain a sustained, altered transcriptional state following injury.

**Figure 1:**
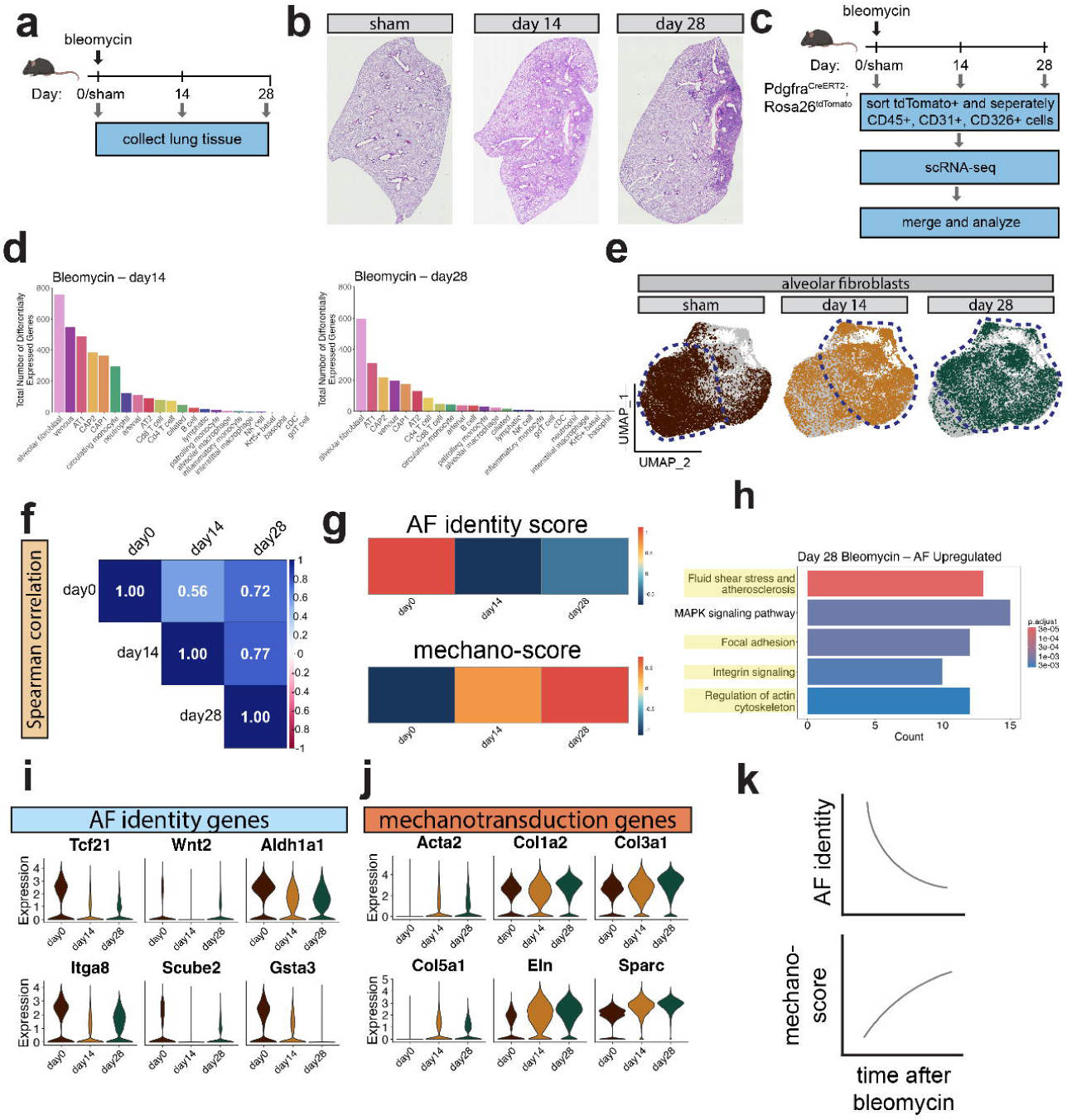
Persistent imbalance of alveolar fibroblast identity after acute lung injury. (a) experimental schematic. (b) H&E-stained mouse lung sections of left lobes at defined timepoints after intratracheal administration of bleomycin. (c) experimental schematic. (d) total number of differentially expressed genes at day 14, and day 28, relative to sham. Cells are ranked by the highest to lowest number of differentially expressed genes at each timepoint. (e) UMAP representation of scRNA-seq data from lineage-traced alveolar fibroblasts after bleomycin. Data represent an integrated dataset. Cells at each timepoint are highlighted within the full dataset and outlined by dotted lines. (f) Spearman correlation analysis, statistically comparing variable genes at each timepoint (sham, day 14, and day 28). (g) heatmaps representing alveolar fibroblast (AF) identity score, and mechano-score over time after bleomycin. (h) Pathway analysis depicting the top 5 terms which were enriched in alveolar fibroblasts at day 28 after bleomycin, relative to cells from sham animals. Terms are ranked by adjusted p-value. (i, j) Violin plots showing expression profiles of representative AF identity and mechanotransduction related genes after bleomycin. (k) summary schematic.

To quantitatively assess transcriptional similarity across time after injury, we performed Spearman rank-order correlation analysis. Although alveolar fibroblasts exhibited a partial transcriptional rebound by day 28, these cells remained transcriptionally distinct and divergent compared to alveolar fibroblasts from healthy animals (Fig. 1f). To determine whether these differences reflected loss of their identity, we generated a module score based on 15 genes selectively enriched in alveolar fibroblasts relative to other mesenchymal and non-mesenchymal lung cell types (Supp. Table 1). Consistent with our Spearman correlation analysis, we observed a persistent, non-resolving reduction in alveolar fibroblast identity after bleomycin (Fig. 1g), accompanied by repression of key genes uniquely expressed in alveolar fibroblasts (Fig. 1i). To determine whether this loss of identity was associated with activation of other pathways, we performed pathway analysis on genes upregulated in alveolar fibroblasts at day 28 after bleomycin. When ranked by adjusted p value, four out of the top five pathways were related to mechanotransduction, including fluid shear stress, focal adhesion, integrin signaling, and regulation of the actin cytoskeleton (Fig. 1h). To further assess the transient mechano-response of alveolar fibroblasts, we generated a “mechano-score” using genes from the KEGG pathways listed above. This analysis revealed a marked increase in mechano-signaling at day 14 after bleomycin that persisted through day 28 (Fig. 1g). This increased mechano-score was associated with sustained upregulation of key mechanotransduction-related genes (e.g. Acta2 and Col1a2) (Fig. 1j). Together, these findings suggest that even as the lung enters the resolution phase of its injury response, alveolar fibroblasts remain in a persistent state of mechanical activation that correlates with a loss of their identity (Fig. 1k).

To determine whether this loss of identity was specific to alveolar fibroblasts or also occurred in other lung cell types, we analyzed our companion scRNA-seq dataset generated from FACS-enriched CD45+ immune, CD31+ endothelial, and CD326+ epithelial cells (Supp Fig 1a). We generated identity module scores for endothelial, epithelial, and immune cell populations using the top 15 genes enriched in each cell type (Supp Table 2). Although several cell types, including AT1 cells and CAP2 endothelial cells showed a stark reduction in their identity after injury, these populations recovered by day 28 after bleomycin (Supp Fig 1b). In stark contrast, alveolar fibroblasts uniquely exhibited a persistent reduction in identity together with sustained elevation of mechano-signaling (Fig. 1g). These data indicate that, although loss of cell identity is a common early response to lung injury, alveolar fibroblasts are distinguished from the response in other cells in the lung by a failure to fully normalize their resting transcriptional state and identity following injury resolution.

### Partial pneumonectomy induces a reversible reduction in alveolar fibroblast identity

Although bleomycin injury partially resolves histologically over time, persistent injury-induced signals can remain in the lung, such as inflammatory cues^27^, raising the question of whether the observed loss of alveolar fibroblast identity reflects an intrinsic response to altered alveolar mechanics or continued exposure to injury-associated signals. To distinguish between these two possibilities, we utilized the partial pneumonectomy (PNX) model to uncouple mechanical cues from cytotoxic injury. In rodents and other small mammals, surgical resection of the left lung lobe triggers compensatory alveolar growth in the remaining lung tissue to restore gas exchange capacity^28–31^. Immediately following resection, PNX induces a transient increase in alveolar strain, followed by neo-alveolarization and regeneration of functional alveolar units in the remnant lung (Fig. 2a)^12,32^. Thus, this model allows for the direct examination of the relationship between in vivo alveolar mechanics and alveolar cell identity in the absence of cytotoxic injury.

**Figure 2:**
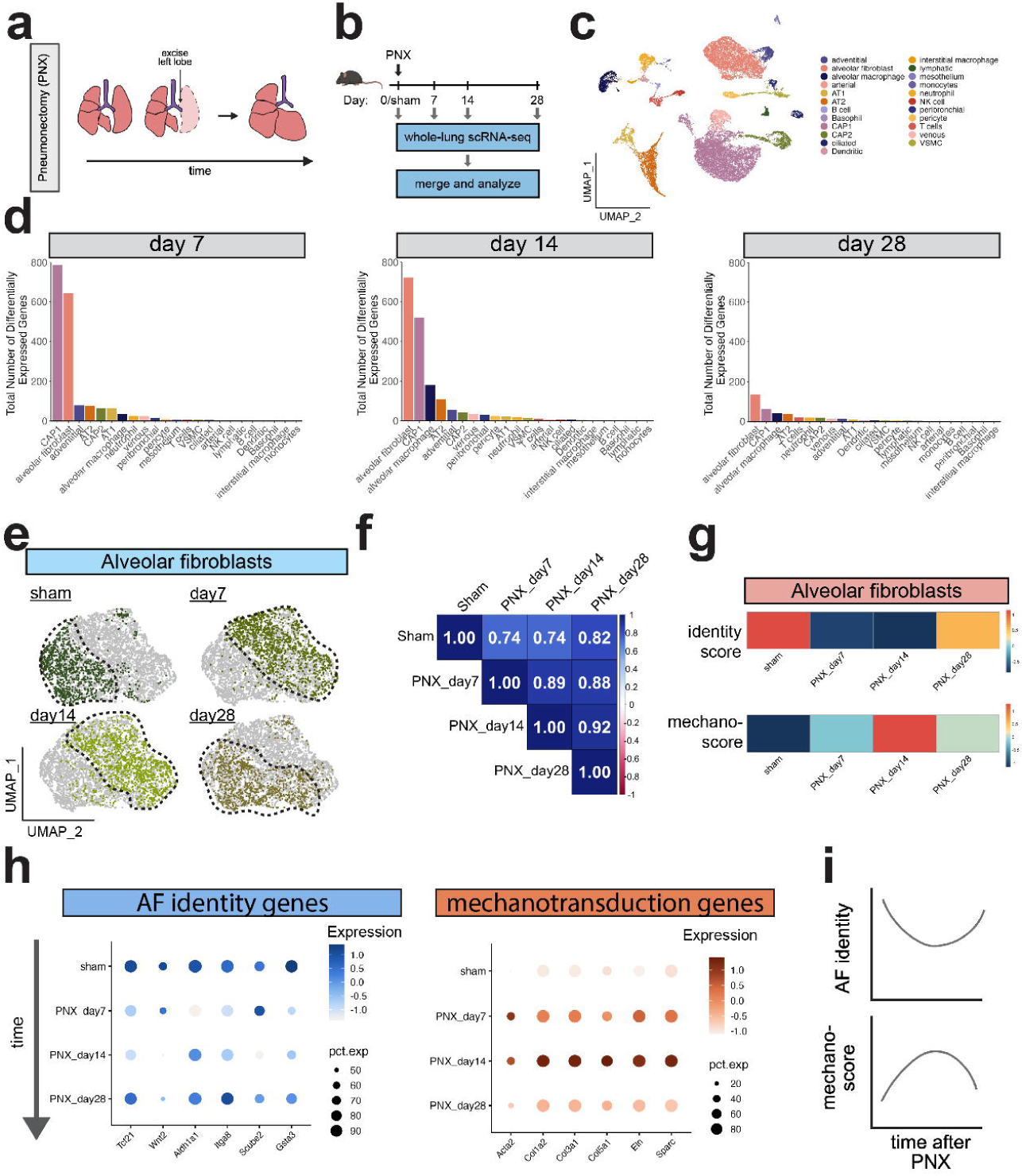
Reversible loss of alveolar fibroblast identity after partial-pneumonectomy. (a) summary schematic depicting partial-pneumonectomy (PNX) model. (b) experimental schematic. (c) UMAP representation of scRNA-seq data. Data represent a merged dataset of all timepoints (sham, day 7, day 14, and day 28). (d) total number of differentially expressed genes at day 7, day 14, and day 28, relative to sham. Cells are ranked by the highest to lowest number of differentially expressed genes at each timepoint. (e) UMAP representation of scRNA-seq data from alveolar fibroblasts after PNX. Data represent an integrated dataset. Cells at each timepoint are highlighted within the full dataset. (f) Spearman correlation analysis, statistically comparing variable genes at each timepoint (sham, day 7, day 14, and day 28) within alveolar fibroblasts. (g) heatmaps representing alveolar fibroblast (AF) identity score, and mechano-score over time after PNX. (h) Dot plots showing expression profiles of representative AF identity and mechanotransduction related genes after partial-pneumonectomy. (i) summary schematic.

To test whether changes in alveolar mechanics are sufficient to modulate alveolar fibroblast identity independent of cytotoxic injury, we performed temporal scRNA-seq after PNX, isolating the right lobes for analysis at days 7, 14, and 28 after resection (Fig. 2b). After clustering, analyzing, and annotating the merged dataset (Fig. 2c), we quantified the total number of differentially expressed genes in each cell type at 7, 14, and 28 days post-PNX compared with sham, providing an unbiased view of how PNX, and the associated changes in mechanical strain, affect all lung cell populations. From this analysis, we identified that most significant changes in alveolar cells occurred at days 7 and 14 post PNX, most of which normalized by day 28 (Fig. 2d). At each timepoint, alveolar fibroblasts exhibited robust transcriptional changes, with notable transcriptional changes also occurring in CAP1 endothelial cells, alveolar macrophages, and alveolar type 2 cells (AT2s) (Fig. 2d). To assess in an unbiased manner how each cell type is responding to the increase in alveolar mechanics, we applied the same mechano-score used in our bleomycin dataset (Fig. 1), for each cell within the PNX dataset (Supp. Fig. 3a). We then ranked each cell type by the strongest increase in mechano-score at day 14 relative to sham cells. We identified alveolar fibroblasts exhibiting the strongest increase in mechano-score relative to all other cell types in the lung, with notable changes also occurring in adventitial fibroblasts, pericytes, and vascular smooth muscle cells (Supp Fig 3b). We then subsetted and re-clustered alveolar fibroblasts and observed distinct UMAP localization driven by timepoint (Fig. 2e), which appeared to resolve by day 28, in contrast to bleomycin (Fig. 1e). Consistent with these findings, we found a robust reduction in Spearman correlation coefficients at day 7 and 14, which progressively recovered by day 28 (Fig. 2f). To assess whether these transcriptional changes were in part due to a loss of identity, we applied the same alveolar fibroblast identity module score from our bleomycin scRNA-seq analysis (Fig. 1g) and identified a robust, but transient, reduction in alveolar fibroblast identity which correlated with an increase in mechano-signaling (Fig. 2g). This loss of identity, and gain of mechano-signaling, correlated with a transient reduction in genes uniquely expressed in alveolar fibroblasts, and upregulation of mechano-signaling genes (Fig. 2h). Thus, in contrast to bleomycin injury, the loss of alveolar fibroblast identity observed after PNX is reversible, suggesting that changes in alveolar tissue mechanics transiently modulate fibroblast identity without permanently destabilizing alveolar fibroblast identity and function (Fig. 2i).

### Loss of respiratory-mediated alveolar stretch stabilizes alveolar fibroblast identity

To further examine the relationship between alveolar stretch and stabilization of alveolar fibroblast identity, we employed the bronchial ligation model, in which the left main bronchus is clipped while the pulmonary vasculature remains intact. This procedure restricts expansion of the ligated lung lobe while maintaining vascular perfusion, thereby reducing respiratory-mediated alveolar stretch (Fig. 3a).

**Figure 3:**
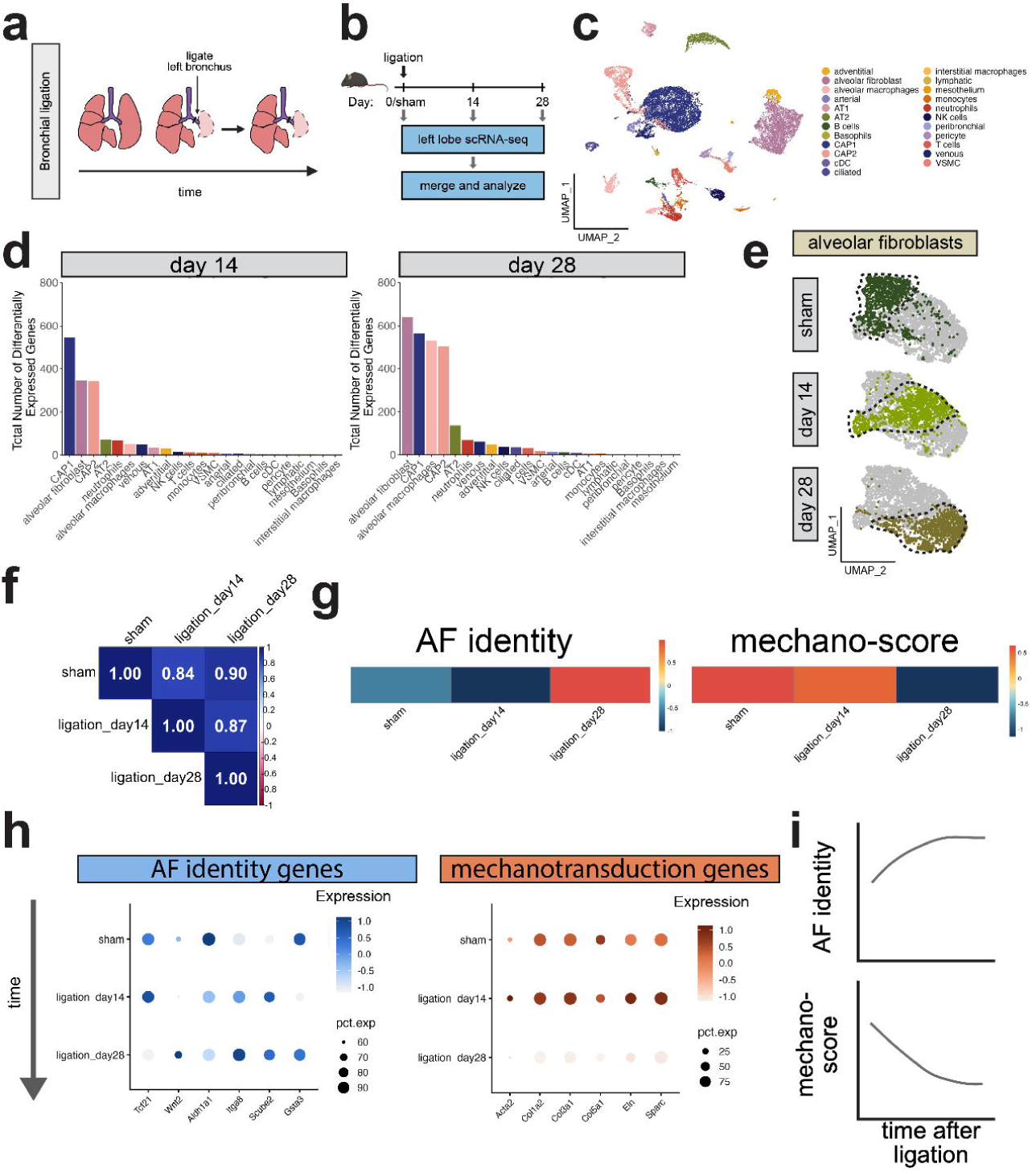
Loss of respiratory stretch stabilizes alveolar fibroblast identity. (a) summary schematic depicting bronchial ligation model. (b) experimental schematic. (c) UMAP representation of scRNA-seq data. Data represent a merged dataset of all timepoints (sham, day 14, and day 28). (d) total number of differentially expressed genes at day 14 and day 28 relative to sham. Cells are ranked by the highest to lowest number of differentially expressed genes at each timepoint. (e) UMAP representation of scRNA-seq data from alveolar fibroblasts after bronchial ligation. Data represent an integrated dataset. Cells at each timepoint are highlighted within the full dataset. (f) Spearman correlation analysis, statistically comparing variable genes at each timepoint (sham, day 14, and day 28) within alveolar fibroblasts. (g) heatmaps representing alveolar fibroblast (AF) identity score, and mechano-score over time after bronchial ligation. (h) Dot plots showing expression profiles of representative AF identity and mechanotransduction related genes after bronchial ligation. (i) summary schematic.

We ligated the left main bronchus and performed scRNA-seq on ligated left lung lobes at 14 and 28 days post-surgery (Fig. 3). After clustering and annotating the merged dataset (Fig. 3c), we quantified the total number of differentially expressed genes in each cell type at days 14 and 28 relative to sham controls. This analysis revealed that alveolar fibroblasts were among the most transcriptionally responsive cell types in this model (Fig. 3d). Additional transcriptional responses were observed in CAP1 and CAP2 endothelial cells, alveolar macrophages, and AT2 cells (Fig. 3d), likely reflecting the AT1-to-AT2 reprogramming response previously reported in this model^10^. In contrast to the transient transcriptional changes observed following PNX, the transcriptional remodeling observed after bronchial ligation persisted across both timepoints, consistent with the sustained loss of alveolar stretch in this model.

To systematically evaluate how mechanical signaling changes across cell types after bronchial ligation, we applied the same mechano-score used in the PNX and bleomycin datasets to each cell within the ligation dataset. Ranking cell types by their change in mechano-score at day 28 relative to sham revealed that AT1 and AT2 epithelial cells exhibited the most pronounced reductions in mechanotransduction, consistent with prior observations that the alveolar epithelium is highly sensitive to loss of respiratory-mediated stretch (Supp Fig. 4)^10^. Importantly, alveolar fibroblasts also exhibited a robust reduction in mechano-score following bronchial ligation (Supp Fig 4).

To further examine fibroblast-specific transcriptional dynamics, we subsetted and re-clustered alveolar fibroblasts and observed progressive transcriptional shifts across time following ligation (Fig. 3e). Spearman correlation analysis confirmed a gradual divergence from the sham transcriptional state that persisted through day 28 (Fig. 3f). To determine how these transcriptional changes affected fibroblast identity, we applied the same alveolar fibroblast identity module score used in our previous analyses. In contrast to the persistent loss of identity observed in bleomycin injury and the transient reduction observed after PNX, bronchial ligation resulted in an increase in alveolar fibroblast identity (Fig. 3g). This increase was associated with elevated expression of genes uniquely enriched in alveolar fibroblasts and a repression of mechanotransduction-associated genes (Fig. 3h).

Taken together, these data demonstrate that loss of respiratory-mediated alveolar stretch suppresses mechanotransduction signaling while reinforcing the transcriptional identity of alveolar fibroblasts (Fig. 3i). When integrated with our bleomycin and PNX analyses, these results suggest that alveolar fibroblast identity is dynamically regulated by the mechanical environment of the alveolus, with increased tissue strain promoting identity destabilization and reduced mechanical stretch stabilizing the alveolar fibroblast state.

### Alveolar fibroblasts remain stable at homeostasis following disruption of nuclear mechanotransduction

Our data thus far have demonstrated that alveolar fibroblasts are uniquely responsive to changes in their mechanical microenvironment. However, whether transmission of mechanical forces to the nucleus is required for this process remains unknown. To address this, we generated mice in which the LINC complex could be selectively ablated in alveolar fibroblasts through conditional deletion of Sun1 and Sun2, thereby severing the connection between the cytoskeleton and the nuclear envelope (Pdgfra^CreERT2^ x Sun1^fl/fl^ x Sun2^fl/fl^ x Rosa26^LSL-tdTomato^, herein referred to as Sun1/2^Pdgfra-KO^) (Fig. 4a). Following tamoxifen administration, we assessed recombination efficiency using qRT-PCR and immunohistochemistry and observed dramatic loss of both Sun1 and Sun2 expression in Pdgfra+ cells (Fig. 4b). Consistent with disruption of mechanotransduction signaling, we also observed reduced expression of the mechanoresponsive genes Ccn1 and Ccn2 (Fig. 4b). Despite disruption of the LINC complex, alveolar fibroblasts persisted for at least six months following Sun1 and Sun2 deletion, indicating that the LINC complex is not required for fibroblast homeostasis in vivo (Fig. 4c). Consistent with these findings, LINC disruption also did not affect fibroblast survival, as apoptosis rates were similar between wildtype and Sun1/2^Pdgfra-KO^ alveolar fibroblasts (Supp Fig. 5b). To determine whether LINC disruption altered nuclear morphology, we quantified cross-sectional nuclear area and observed no significant difference between Sun1/2^Pdgfra-KO^ and wildtype Pdgfra+ fibroblasts (Fig. 4d, Supp Fig. 5a).

**Figure 4:**
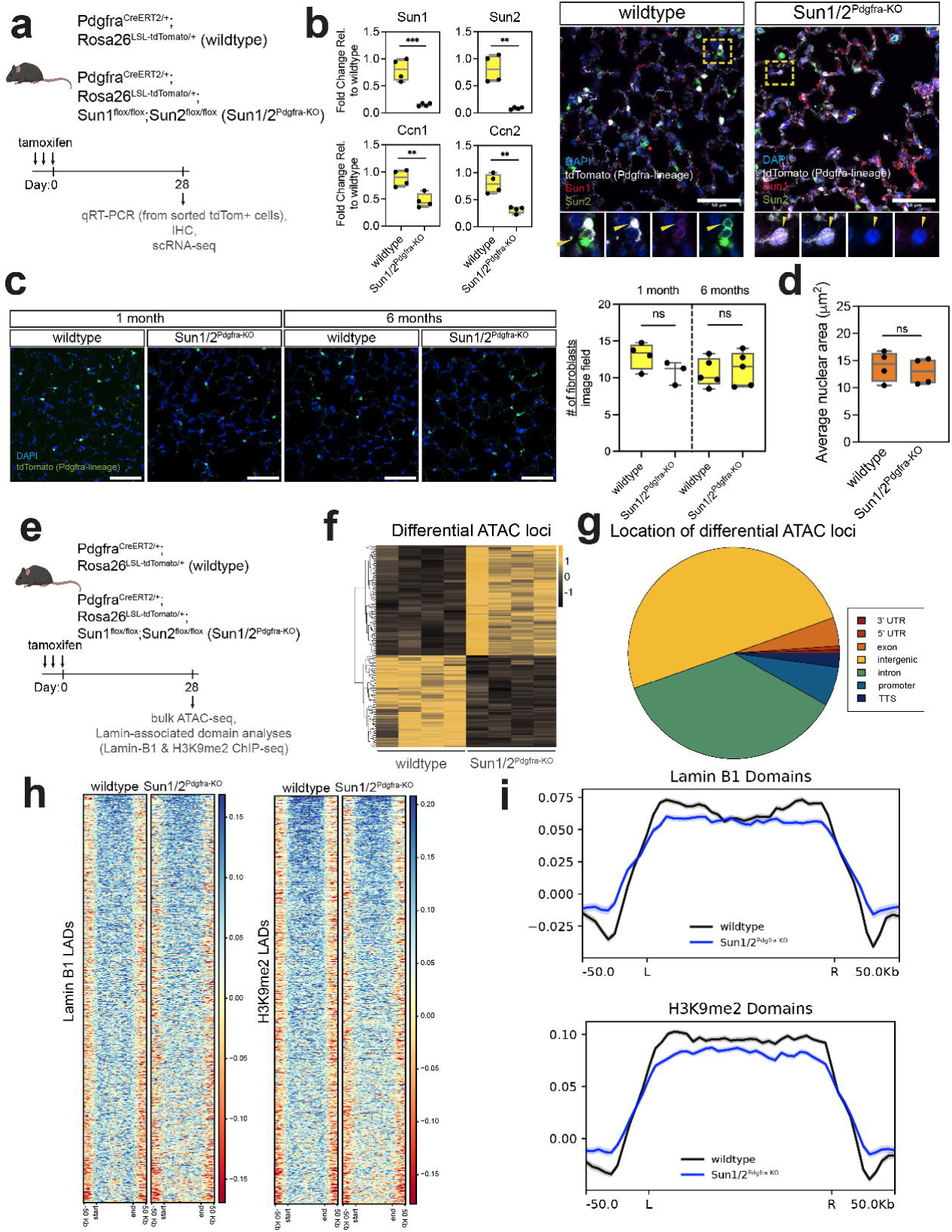
Alveolar fibroblasts remain stable following loss of LINC complex function. (a) experimental schematic. (b) qRT-PCR and IHC of wildtype and Sun1/2^Pdgfra-KO^ lung fibroblasts confirming robust depletion of Sun1 and Sun2 in alveolar Pdgfra+ lung fibroblasts. (c) IHC and quantification demonstrating persistence of alveolar Pdgfra+ lung fibroblasts (tdTomato+ cells, shown in green) over time after Sun1 and Sun2 deletion. (d) average nuclear area, assessed by nuclear lamin A/C staining comparing wildtype and Sun1/2^Pdgfra-KO^ lung fibroblasts. (e) experimental schematic. (f) heatmap of the differentially accessible genomic loci comparing wildtype and Sun1/2^Pdgfra-KO^ lung fibroblasts. (g) genomic distribution of the differential accessibility sites in panel (f). (h) Heatmap depicting LAD signal intensity of both Lamin B1 and H3K9me2 domains between wildtype and Sun1/2^Pdgfra-KO^ fibroblasts. (i) Signal profile of Lamin B1 and H3K9me2 across LAD domains between wildtype and Sun1/2^Pdgfra-KO^ fibroblasts.

To test whether the LINC complex regulates chromatin structure under homeostatic conditions, we isolated alveolar fibroblasts from wildtype and Sun1/2^Pdgfra-KO^ mice and performed bulk ATAC-seq alongside Lamin B1 and H3K9me2 ChIP-seq to assess nuclear lamina– chromatin interactions, focusing on lamina-associated heterochromatin domains (LADs) (Fig. 4e). ATAC-seq analysis revealed that depletion of the LINC complex altered chromatin accessibility at a limited number of loci, with increased accessibility at 100 loci and decreased accessibility at 84 loci (Fig. 4f, g). Although biological replicates were highly consistent based on PCA and hierarchical clustering analyses (Supp Fig. 6a, b), these loci were not enriched for genes associated with canonical mechanotransduction pathways nor alveolar fibroblast identity (Supp Fig. 6c, Supp Table 3), suggesting that LINC disruption has only modest effects on chromatin accessibility under homeostatic conditions. To determine whether LINC depletion alters nuclear lamina organization, we next analyzed Lamin B1 and H3K9me2 ChIP-seq datasets to evaluate LAD architecture. Surprisingly, this analysis revealed strong correlation between genotypes, indicating minimal changes in LAD formation between wildtype and Sun1/2^Pdgfra-KO^ fibroblasts (Fig. 4h, Supp Fig. 7). Quantitative analyses further confirmed no significant differences in LAD abundance, number, signal intensity, or domain width between wildtype and Sun1/2^Pdgfra-KO^ fibroblasts (Fig. 4i).

Together, these findings demonstrate that disruption of the LINC complex at homeostasis has minimal effects on alveolar fibroblast stability, chromatin accessibility, and lamina-associated domain architecture, indicating that nuclear mechanotransduction is dispensable for maintenance of the homeostatic fibroblast state. These results are also consistent with the lung normally existing in a quiescent state and suggest that LINC-mediated force transmission may shape fibroblast responses to injury.

### Disruption of nuclear mechanotransduction promotes lung regeneration

We next examined whether severing cytoskeletal-nuclear force transmission alters the regenerative response following lung injury. We performed bleomycin-induced lung injury in both Sun1/2^Pdgfra-KO^ and wildtype mice and collected lungs at 14 and 28 days post injury for analysis (Fig. 5a). To measure bleomycin-induced lung tissue damage and scarring, we performed picrosirius red (PSR) staining and quantification. While we observed no significant difference in PSR+ lung area between Sun1/2^Pdgfra-KO^ and wildtype mice at 14 days, we detected a significant reduction in PSR+ area in Sun1/2^Pdgfra-KO^ animals by 28 days post injury (Fig. 5b, c). A similar trend was observed from quantitative H&E analyses, where we observed a reduction in severely damaged lung tissue in Sun1/2^Pdgfra-KO^ mice at 28 days post injury (Fig. 5d, e). Given this histological evidence of improved tissue regeneration, we examined whether ablation of the LINC complex disrupted the potential of alveolar fibroblasts to proliferate or differentiate into myofibroblasts. Interestingly, we found that deletion of the LINC complex did not alter the capacity of alveolar fibroblasts to proliferate or differentiate into myofibroblasts after injury (Supp. Fig. 8a, b). Our previous work has shown that alterations in alveolar fibroblast function can induce proliferation changes in AT2 cells^5^, a cardinal feature in lung alveolar regeneration. Loss of the LINC complex in Pdgfra+ cells resulted in a significant increase in proliferating AT2 cells at 14 days post-injury relative to wildtype controls (Fig. 5f, g). Furthermore, loss of the LINC complex in alveolar fibroblasts nearly abolished the dysplastic Krt5+ cell response at 28 days post injury, while control animals exhibited obvious dysplasia (Fig. 5h). Taken together, these findings indicate that ablation of the LINC complex promotes pro-regenerative signaling and enhances lung repair.

**Figure 5:**
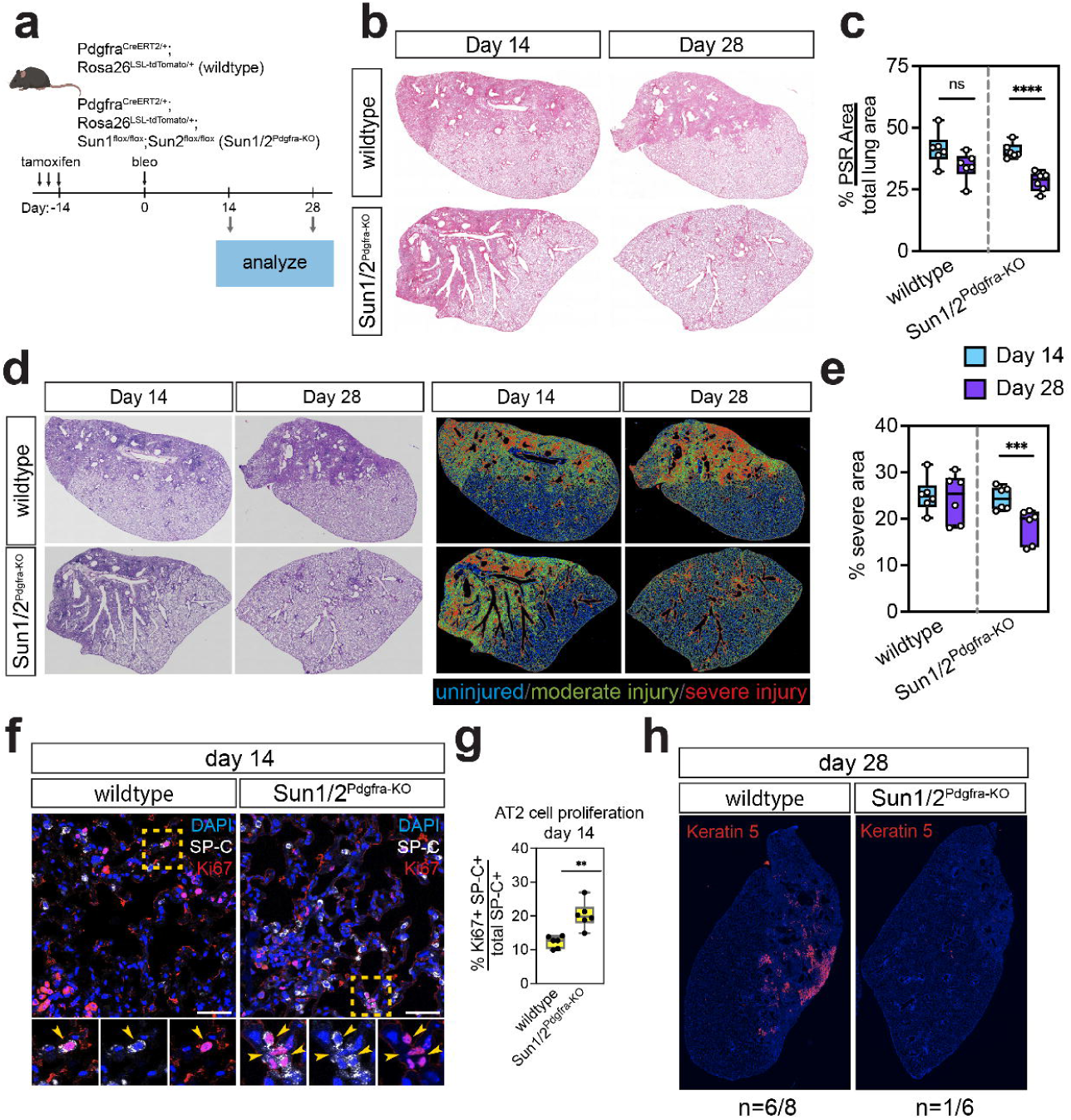
LINC depletion in alveolar fibroblasts enhances alveolar regeneration after lung injury. (a) Schematic depicting approach to assess the role of Sun1 and Sun2 in lung repair and regeneration. (b) Representative picrosirius red staining (PSR) of the left lobes after Sun1/2 deletion in Pdgfra+ cells after bleomycin-induced lung injury and (c) quantification of PSR+ area as a percentage of total lung area. (d) Representative hematoxylin and eosin (H&E) staining of the left lobes after Sun1/2 deletion in Pdgfra+ cells after bleomycin-induced lung injury and (e) quantification of injury severities as a percentage of total lung area. (f) representative IHC images showing an increase in proliferation of AT2 cells after Sun1 and Sun2 deletion in Pdgfra+ cells and (g) quantification. Scale bar is 50 um. Each dot represents data obtained from a single animal (biological replicate). **P<0.01 evaluated by unpaired t-test. (h) representative IHC images showing a decrease in dysplastic Krt5+ basal cells after Sun1 and Sun2 deletion in Pdgfra+ cells.

### Loss of LINC complex function resolves fibroblast identity after injury

Given that disruption of the LINC complex enhanced lung regeneration after injury, we next asked whether this effect was mediated by restoration of alveolar fibroblast identity. To test this, we performed scRNA-seq at 14 and 28 days after bleomycin injury, comparing isolated Pdgfra+ cells from Sun1/2^Pdgfra-KO^ animals with previously generated wildtype control animals (Fig. 6a). Cells were analyzed based on genotype and timepoint, revealing clear transcriptional separation between fibroblasts from Sun1/2^Pdgfra-KO^ and wildtype animals (Fig. 6b). Notably, at 14 days post-injury, fibroblasts from both genotypes converged transcriptionally, indicating a shared early injury response. Consistent with our histological analyses (Supp Fig 8), Sun1/2^Pdgfra-KO^ alveolar fibroblasts retained the ability to proliferate (Mki67+, Top2a+) and adopt a myofibroblast phenotype (Acta2+, Tagln+) following injury (Supp. Fig. 8c). In addition to these canonical injury-associated states, we identified an inflammatory fibroblast cluster in Sun1/2^Pdgfra-KO^ animals marked by Saa3 and Lcn2 expression, consistent with inflammatory fibroblast states reported in other lung injury models^5,6,33,34^. However, by 28 days post-injury, fibroblasts from Sun1/2^Pdgfra-KO^ animals largely returned to a transcriptional state resembling their uninjured state, whereas wildtype injured fibroblasts remained in a more activated state (Fig. 6b).

**Figure 6:**
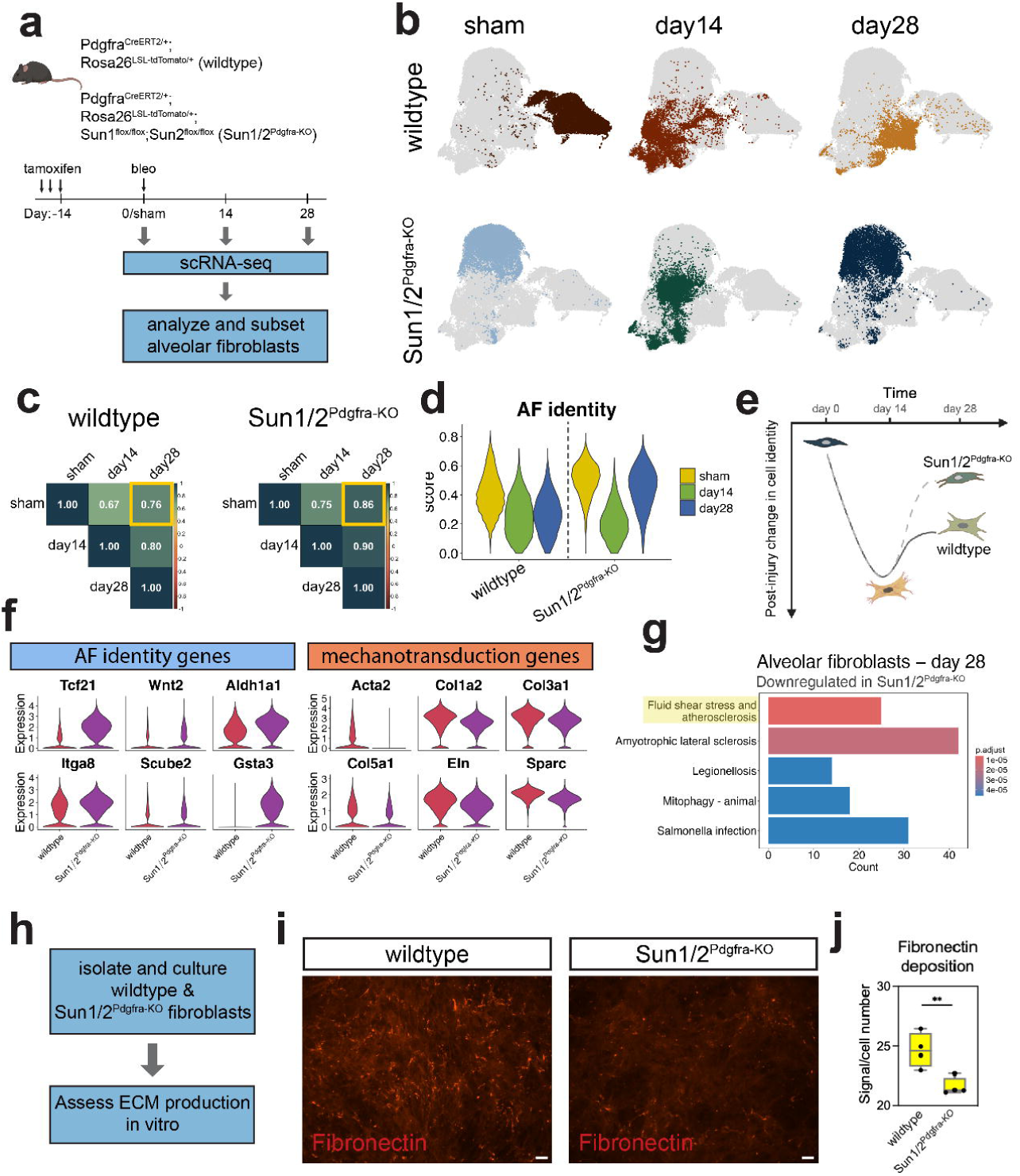
LINC depletion normalizes alveolar fibroblast identity after lung injury. (a) Schematic depicting approach to assess the role of Sun1 and Sun2 in lung repair and regeneration. (b) UMAP representation of scRNA-seq data of alveolar Pdgfra+ fibroblasts highlighting UMAP localization of cells at each timepoint (sham, day 14, and day 28) vs genotype (wildtype vs Sun1/2^Pdgfra-KO^). (c) Correlation plot showing Spearman correlation coefficients of wildtype and Sun1/2^Pdgfra-KO^ alveolar fibroblasts over time after bleomycin-induced injury. (d) violin plots demonstrating a rescue of alveolar fibroblast identity in Sun1/2^Pdgfra-KO^ animals at day 28 after bleomycin. (e) schematic demonstrating a rescue of alveolar fibroblast identity after bleomycin following Sun1 and Sun2 deletion. (f) Violin plots demonstrating a rescue of alveolar fibroblast identity genes, and a reduction in mechanotransduction related genes in Sun1/2^Pdgfra-KO^ cells relative to wildtype cells. (g) Pathway analysis of the gene expression programs which are downregulated in alveolar Sun1/2^Pdgfra-KO^ cells at 28 days post bleomycin. (h) schematic depicting approach to assess ECM production in wildtype vs Sun1/2^Pdgfra-KO^ fibroblasts in vitro. (i) representative immunofluorescence images and (j) quantification of deposited Fibronectin between wildtype and Sun1/2^Pdgfra-KO^ lung fibroblasts after 6 days in culture. Scale bar represents 50 um. Each dot represents data obtained from a single mouse (biological replicate). **P<0.01 evaluated by unpaired t-test.

To quantify these transcriptional changes, we performed Spearman correlation analysis comparing injured fibroblasts to their homeostatic counterparts. Consistent with the clustering results, alveolar fibroblasts from Sun1/2^Pdgfra-KO^ animals at 28 days post-injury exhibited a stronger correlation with homeostatic fibroblasts (ρ = 0.90) than injured wildtype fibroblasts (ρ = 0.79) (Fig. 6c). We next examined whether this normalization of transcriptional state was accompanied by restoration of alveolar fibroblast identity. Application of the same alveolar fibroblast identity module score used in our earlier analyses revealed that fibroblasts from both genotypes initially exhibited reduced alveolar identity at 14 days post-injury (Fig. 6d). However, while wildtype fibroblasts exhibited persistent loss of alveolar identity at later timepoints, Sun1/2^Pdgfra-KO^ fibroblasts normalized their identity by 28 days post-injury back to their inherent homeostatic state (Fig. 6d, e). This recovery was accompanied by increased expression of genes uniquely enriched in alveolar fibroblasts and reduced expression of mechanotransduction-associated genes (Fig. 6f, Supp Fig. 8d). Improved recovery of key signaling molecules, such as Wnt2 and Fgf7, is a likely mechanism underlying the enhanced epithelial regenerative response noted above (Supp Fig. 8d).

To further determine whether mechanotransduction pathways were reduced following LINC disruption, we performed pathway analysis on genes downregulated in Sun1/2^Pdgfra-KO^ fibroblasts relative to wildtype fibroblasts at 28 days post-bleomycin. This analysis revealed enrichment of mechanotransduction-related pathways, including the fluid shear stress pathway previously identified in our bleomycin dataset in Fig. 1 (Fig. 6g). To determine whether this transcriptional normalization of cell identity was accompanied by changes in core fibroblast functions, we isolated primary fibroblasts from Sun1/2^Pdgfra-KO^ and wildtype animals and measured ECM deposition in vitro (Fig. 6h). Consistent with our transcriptional analyses, Sun1/2^Pdgfra-KO^ fibroblasts exhibited reduced fibronectin deposition relative to wildtype controls (Fig. 6i, j).

Taken together, these data demonstrate that ablation of the LINC complex enables alveolar fibroblasts to exit a persistent mechano-activated injury state and return to a homeostatic transcriptional identity, thereby promoting a regenerative microenvironment that supports functional lung repair.

## Discussion

Alveolar fibroblasts are increasingly recognized as a specialized mesenchymal cell population that supports epithelial regeneration in the distal lung^35^. Although these cells are known to activate after injury^5–7,33^, the mechanisms that preserve or destabilize their cell identity during tissue repair have remained unclear. In this study, we identify alveolar tissue mechanics as a major regulator of alveolar fibroblast identity and show that nuclear mechanotransduction through the LINC complex contributes to the persistence of an injury-associated fibroblast state that limits regenerative repair of the lung. Across multiple in vivo models, we find that increased mechanotransduction is associated with destabilization of alveolar fibroblast identity, whereas loss of mechanotransduction reinforces the alveolar fibroblast state. Moreover, disrupting cytoskeletal-nuclear force transmission through the LINC complex enables fibroblasts to more effectively normalize their homeostatic identity after injury, thereby promoting lung regeneration.

Our findings identify tissue mechanics as a central regulator of alveolar fibroblast identity and position nuclear mechanotransduction through the LINC complex as a key determinant of whether fibroblasts maintain or lose their niche-supportive state. Consistent with this framework, recent work demonstrates that fibroblast cell states can be dynamically regulated by mechanical inputs in non-pulmonary contexts, supporting a broader principle that mechanotransduction governs fibroblast state stability^36,37^. In the lung, we find that increased mechanical stress is associated with destabilization of alveolar fibroblast identity, whereas disruption of force transmission through the LINC complex reinforces the homeostatic state and promotes recovery following injury. Together, these findings support a model in which fibroblast identity is continuously tuned by the physical microenvironment and suggest that failure to appropriately resolve mechanical signaling may underlie the persistence of maladaptive fibroblast states that impair lung regeneration.

Our observations are consistent with prior work demonstrating that mechanical environments can destabilize fibroblast cell states. Several studies have shown that fibroblasts cultured on stiff substrates that mimic fibrotic tissue adopt persistent myofibroblast-like transcriptional programs and exhibit altered nuclear architecture and chromatin organization^13,14^. In parallel, recent work has demonstrated that mechanical tension transmitted to the nucleus can stabilize fibroblast activation by promoting chromatin condensation and persistent transcriptional changes, and that uncoupling cytoskeletal tension from the nucleus prevents stabilization of the activated fibroblast state^37^. Together, these studies suggest that mechanical forces can imprint transcriptional programs in fibroblasts through nuclear mechano-sensing mechanisms. Our results reveal that alveolar mechanics regulate fibroblast identity within the lung and that disruption of nuclear force transmission prevents the persistence of an injury-associated fibroblast state during tissue regeneration.

In summary, we identify alveolar fibroblast identity as a mechanically regulated and injury-sensitive cell state that plays a central role in determining regenerative outcome after lung injury. By integrating comparative in vivo models of altered alveolar mechanics with genetic disruption of cytoskeletal-nuclear force transmission, we show that the LINC complex promotes persistence of an injury-associated fibroblast state, whereas uncoupling mechanical signals from the nucleus facilitates recovery of homeostatic identity and improves lung repair. These findings establish nuclear mechanotransduction as a key regulator of alveolar fibroblast identity and suggest that restoring stromal cell identity may represent a therapeutic strategy to enhance regeneration in fibrotic lung disease.

## Methods

### Animals

All mouse experiments were performed under the protocols approved by the guidance of the University of Pennsylvania Institutional Animal Care and Use Committee. Pdgfra^CreERT2^ and Rosa26^tdTomato^ mouse lines have been previously described^38,39^ (Jax stocks 032770 and 007914, respectively). Sun1-flox and Sun2-flox mice were generated by crossing the Sun1^tm1a^ (C57BL/6N-Sun1^tm1a(EUCOMM)Wtsi^/ CipheOrl, purchased from InfraFrontier/EMMA, stock #09532) and Sun2^tm1a^ (C57BL/6N-A^tm1Brd^ Sun2^tm1a(EUCOMM)Hmgu^/BayMmucd, purchased from MMRRC, stock #037788-UCD) alleles with FLPo deleter mice (Jax stock 012930)^40^ to remove the LacZ cassettes. All experiments were performed on 8-12 week old mice that were maintained on a mixed C57BL/6 and CD1 background. Both male and female mice were used in all conditions.

### Tamoxifen administration

For Cre recombinase induction, tamoxifen (Millipore Sigma) was dissolved in corn oil and ethanol mixture (90%/10%, v/v) (Millipore Sigma) to produce a stock solution with a concentration of 20 mg/mL. Pdgfra^CreERT2^ mice were administered tamoxifen via intraperitoneal injections at a dose of 200 mg/kg for 3 consecutive days.

### Bleomycin lung injury

Two weeks after tamoxifen induction, 3 U/kg bleomycin (Teva), diluted in PBS (Thermo Fisher Scientific), was administered intratracheally to anesthetized mice. Control mice received sterile PBS.

### Unilateral lung ligation

Mice were anesthetized with isoflurane and intubated for ventilation using a MiniVent ventilator (Harvard Apparatus) as previously described^10^. A 2 cm skin incision was made along the left lateral thorax, followed by a 1 centimeter incision through the fifth left intercostal space. The left main bronchus was then occluded using a Micro-clip (Horizon). The ligated left lung lobe was analyzed 14 and 28 days after the procedure.

### Partial pneumonectomy

After exposing the left thoracic cavity as described above, the left main bronchus and pulmonary vasculature were ligated with sterile thread, followed by resection of the left lung lobe. Right lung lobes were analyzed on days 7, 14, and 28 after the procedure.

### Histology and IHC

Mice were euthanized by CO_2_ inhalation and the lungs were perfused with ice-cold PBS through the right ventricle. The lungs were then inflated with 2% PFA (Thermo Fisher Scientific) at a constant pressure of 25 cm H_2_O and were fixed overnight at 4 °C. Tissue was then dehydrated in a series of ethanol concentration gradients, paraffin embedded, and 6um thick sections were cut. Hematoxylin and eosin staining was performed as previously described^5^. For IHC, the following antibodies were used on paraffin sections: RFP (goat, Origene, cat# AB8181-200), RFP (rabbit, Rockland, cat# 600-401-379), SMA (goat, Novus Biologicals, cat# NB300-978), SMA (rabbit, Abcam, cat# ab5694), Ki67 (mouse, BD Biosciences, cat# 550609), SP-C (rabbit, Millipore-Sigma, cat# AB3786), DC-Lamp (Novus, cat# DDX0191P-100), Cleaved Caspase-3 (R&D Systems, cat# MAB835), Keratin5 (rabbit, Abcam, cat# ab52635), Sun1 (mouse, Millipore-Sigma, cat# MABT892), Sun2 (rabbit, Abcam, cat# ab124916).

### Imaging and image analysis

Fluorescent images were acquired at 40x using z-stacks on an LSM 710 laser scanning confocal microscope (Zeiss) and Stellaris 5 laser scanning confocal microscope (Leica). Cell counting analysis was completed within FIJI. Picrosirius Red (PSR) and H&E stained lung sections were tile-scanned using a 4x objective on the Eclipse Ni series upright microscope (Nikon). For the PSR scanning, all images were acquired using the same light and camera settings. Whole lobe immunofluorescence tile scans in Figure 5e were acquired using a 4x objective on the Nikon Eclipse Ni series upright microscope. Epithelial dysplasia, defined by expansion of Krt5+ epithelium into the alveolar parenchyma, was scored in lungs exhibiting the phenotype shown in Figure 5.

### H&E quantification

To assess injury severity based on H&E staining, we used a previously published algorithm^5,41^. Briefly, we used the EBImage package with R. We then imported images and converted RGB pixel intensity to a matrix and then applied K-means clustering to segment out, in an unbiased manner, 4 different regions of injury severity: normal, moderate, severe, and background. These regions correspond to colors which are coded as follows: blue, green, red, and black. Using the area of the lobe outlined in H&E, the percent of each injured zone (normal, moderate, and severe) were then calculated as the percent of total lung lobe area. This R script has been versioned and can be accessed on the Morrisey Lab Github page (https://github.com/Morriseylab).

### Picrosirius red staining and quantification

Picrosirius red (PSR) staining was performed according to manufacturer’s protocol (Polysciences, cat # 24901). PSR stained lung sections were imaged as described above. Tile stitched images were then analyzed within Fiji.

### Lung tissue digestion and FACS

Lungs were harvested and digested into single cell suspensions using collagenase I (Thermo Fisher, cat# 17100017), dispase (Corning, cat# 354235), and DNase I (Millipore Sigma, cat # 4716728001), as previously described^5,41^. Red blood cells were removed with ACK lysis buffer (Quality Biological, cat# 118-156-101) and then the cell suspension was stained with antibodies diluted in ice-cold FACS buffer [PBS, 25mM HEPES (Thermo Fisher Scientific, cat# 15630080), 2mM EDTA (Invitrogen, cat# AM9260G), and 2% Fetal Bovine Serum (FBS) (Corning, cat# 35-015-CV)]. The following antibodies were used for flow cytometry and cell sorting: CD45-APC (1:200, ThermoFisher, clone: 30-F11, cat. 17-0451-83), CD31-APC (1:100, ThermoFisher, clone: 390, cat. 17-0311-82), and CD326(aka EpCAM)-APC (1:300, ThermoFisher, clone: G8.8, cat. 17-5791-82). Cells were stained with antibodies (diluted in FACS buffer) for 10 minutes on ice. Cells were then spun down at 300g for 5 minutes and resuspended in ice cold FACS buffer. Cells were then stained with anti-APC microbeads (1:100, Miltenyi Biotec, cat# 130-090-855) for 10 minutes on ice. Cells were then spun down at 300g for 5 minutes and resuspended in ice-cold PBS + 0.5% BSA (Jackson Immunoresearch, cat# 001-000-162). Cells were then passed through a magnetic LS Column (Miltenyi Biotech, cat#130-042-40). LS Columns were washed with 10mL of ice-cold PBS + 0.5% BSA. After passing through LS Columns, cells were then spun down at 300g for 5 minutes and resuspended in 400 uL of ice-cold FACS buffer. DRAQ7 (BD Biosciences, cat# 564904) was added to the cell suspension (1:100) prior to sorting. The following gating strategy was used to sort tdTomato+ Pdgfra-lineage traced fibroblasts: SSC vs FSC, Trigger Pulse width vs FSC, SSC vs DRAQ7, tdTomato (Pdgfra+ cells) vs APC. All fibroblasts (tdTomato+) were sorted using a 100μm sized nozzle into ice-cold FACS buffer using the cell sorter FACSJazz (BD Biosciences).

### scRNA-seq and analysis

For the whole-lung scRNA-seq data in Figs. 1 and 2, mouse lungs were harvested and digested as described above and previously^5^. Cell suspensions were incubated with anti-CD45 mouse microbeads (Miltenyi Biotec, cat# 130-052-301) for 30 minutes on ice. Cells were then passed through a magnetic LS column to deplete the cell suspension of CD45+ immune cells. CD45+ immune cells were then later recovered and spiked back into the CD45-depleted cell suspension at a concentration of 20% of the total cell population. The resultant cell mixture was then loaded onto the 10x Chromium controller (10x Genomics). For the Pdgfra-lineage trace fibroblasts scRNA-seq, cells were sorted using the tdTomato reporter as described previously^5^.

All samples were loaded to aim for a recovery of 10,000 cells, and libraries were prepped according to the manufacturer’s protocol using the Chromium Single Cell 3′ v3.1 chemistry. Libraries were then sequenced on an Illumina Novaseq 6000 instrument and Illumina Novaseq X instrument. The sequenced data was processed by aligning reads and obtaining distinct molecular identifiers (UMIs) using STARsolo (v2.7.9a). The scRNA-seq data was further processed and analyzed using the Seurat v4 package^42^. For the whole-lung scRNA-seq data in Figures 1 and 2, cells were removed if: (i) the number of genes detected was less than 800 or greater than 5000, (ii) if the number of counts were greater than 20,000, (iii) if the percent mitochondrial reads were greater than 20%, and (iv) if the bcds score was greater than 0.8 (R package scExtras v1.0.1). For the scRNA-seq data from lineage traced Pdgfra+ fibroblasts in Figures 5 and 6, cells were removed if: (i) the number of detected genes was less than 500 or greater than 4000, (ii) if the percent mitochondrial reads were greater than 15%, and (iii) if the bcds score was greater than 0.8. For all data, feature (gene) data was scaled in order to remove unwanted sources of variation using the Seurat SCTransform function based on percent mitochondrial reads, and the number of genes, and total reads. Non-linear dimension reduction was performed using uniform manifold approximation and projections (UMAPs) (reduction = “pca” and n.neighbors = 15) and Louvain graph-based clustering algorithms.

Marker genes for each cell type were identified using the FindAllMarkers command using the RNA assay within Seurat. Mesenchymal cells were annotated using previously published canonical marker genes. Epithelial, endothelial, and immune cells were annotated using marker genes and LungMAP labels^43^. Module scores were calculated using UCell. The “mechano-score” module was generated using the unique genes in the KEGG pathway IDs: mmu05418, mmu04510, mmu04518, mmu04810. All gene lists used for module scores are in Supp. Tables 1 and 2.

Spearman correlation analysis was performed in R. Transcription factor activity scores were assigned to cells using the DoRothEA and decoupleR database within R^44–46^. clusterProfiler and DAVID were used for KEGG and GO analyses^47,48^. Visualization of corrplot and transcription factor activity scores were generated using functions within each software package, base R, or ggplot2. UMAPs, violin plots, and dot plots were generated using either Seurat or scCustomize using the RNA assay.

### Bulk ATAC-seq and analysis

Bulk ATAC-seq was performed as previously described following the published protocol^49,50^. DNA fragment size was analyzed using a Fragment Analyzer. Libraries were sequenced to 150 base pairs from both ends using an Illumina NovaSeq. Raw reads were trimmed to 60 base pairs using Trimmomatic^51^, and then aligned to the mm10 genome using Bowtie2^52^. Picard and Samtools were used to generate bam files and to filter out duplicates and mitochondrial reads^53^. Peaks were called using MACS2 using the following options: “--keep-dup all -q 0.01 -no model”^54^. Diffbind was used to identify differentially accessible chromatin loci between wildtype and Sun1/2^Pdgfra-KO^ groups (n = 4 biological replicates/group)^55^. Homer was used to annotate peaks to genomic regions and to the nearest transcriptional start site^56^.

### ChIP-seq and analysis

#### ChIP

Wildtype and Sun1/2^Pdgfra-KO^ were isolated as described above and processed for ChIP as previously described^57^. Wildtype and Sun1/2^Pdgfra-KO^ cells were crosslinked post isolation using methanol-free formaldehyde for 10 min at room temperature with gentle rotation (final 1% v/v). Crosslinking was quenched by addition of glycine for 5 min at room temperature with gentle rotation (final 125mM). Cells were collected by centrifugation (250xg for 5 minutes at room temperature), washed once in 1 x PBS, and collected again by centrifugation. Resulting pellets were flash frozen on dry ice and stored at −80°C. For each ChIP sample, 30µL protein G magnetic beads (Invitrogen #10004D) were washed 3 times in blocking buffer (0.5% BSA in 1x PBS) before beads were resuspended in 250µL blocking buffer plus 2µg antibody (anti-LaminB1 [Abcam #ab16048] or anti-H3K9me2 [Abcam #ab1220]) and rotated at 4°C overnight. Nuclei were isolated from frozen pellets as follows: pellets were resuspended in 10mL cold Lysis Buffer 1 (50mM HEPES-KOH pH7.5, 1mMEDTA, 140mM NaCl, 10% Glycerol, 0.5% NP-40, 0.25% TritonX-100, and protease inhibitor) and rotated at 4°C for 10 min. Resuspended cells were centrifuged at 250xg for 5 min at room temperature and supernatant discarded. Remaining pellet was resuspended in 10mL cold Lysis buffer (220mM NaCl, 10mM Tris-HCl pH 8.0, 1mM EDTA, 0.5mM EGTA, and protease inhibitor) and rotated at 4°C for 10 min. Resuspended cells were centrifuged at 250xg for 5 min at room temperature and supernatant discarded. Nuclei were resuspended in 1mL cold Lysis Buffer 3 (10mM Tris-HCl pH 8.0, 100mM NaCl, 1mM EDTA, 0.5mM EGTA, 0.1% Na-Deoxycholate, and protease inhibitor). Samples were transferred to a pre-chilled 1mL Covaris AFA tube (Covaris #520130) and sonicated using a Covaris S220 sonicator (high cell chromatin shearing for 15 min). TritonX-100 (final 1%) was added to lysates and centrifuged at top speed for 10 min at 4°C. Supernatant was collected, and protein concentration measured via Bradford assay. Antibody-conjugated beads were washed 3 times in blocking buffer and resuspended in 50µL blocking buffer before 500µg protein lysate was added and allowed to rotate overnight at 4°C. For inputs, 50µg protein lysate was aliquoted and stored at −20°C. The next day, antibody-conjugated beads with lysate were washed 5 times for 2 min each at 4°C with rotation in 1mL RIPA buffer (50mM HEPES-KOH pH 7.5, 500mM LiCl, 1mM EDTA, 1% NP-40, 0.7% Na-Deoxycholate). Beads were washed in 1mL Final Wash buffer (1XTE, 50mM NaCl) for 2 min with rotation at room temperature. Lastly, beads were resuspended in 210µL Elution Buffer (50mM Tris-HCl pH 8.0, 10mM EDTA, 1% SDS) and incubated at 65°C for 30 min with agitation. 200µL eluate was removed and all samples were incubated at 65°C with agitation for a minimum of 12 h, but no more than 18 h. 200µL of 1X TE was added the resulting samples followed by addition of RNase (final 0.2mg/mL, Sigma #10109169001) and incubated at 37°C for 2 h. Proteinase K was subsequently added (final 0.2mg/mL, Roche #3115879001) and incubated at 55°C for 2 h. DNA was isolated via phenol:chloroform extraction and resuspended in 10mM Tris-HCl pH8.0. DNA was quantified by Qubit.

#### ChIP-seq

ChIP-seq libraries were prepared using NEBNext Ultra II DNA library prep kit (NEB #E7770L). Samples were prepared with dual unique indices for multiplex sequencing using NEBNext Multiplex Oligos (NEB #E7630). Library quality was determined by BioAnalyzer or 2% agarose gel electrophoresis and quantified by qPCR using NEBNext Library Quant Kit (NEB #E7630S). Libraries were pooled, re-quantified, and single-end sequenced on the Illumina NextSeq1000. Base call (BCL) files were demultiplexed in Illumina BaseSpace Sequence Hub using the sample-specific dual-index combinations provided during library preparation. Demultiplexing was performed with the integrated bcl2fastq pipeline, and the resulting FASTQ files were downloaded for downstream analysis.

Read quality was assessed using FastQC (v0.12.1) and summarized with MultiQC. Adapter trimming, low-complexity read removal, and PCR duplicate removal were performed with fastp (v0.23.2)^58,59^ using default parameters unless otherwise stated. Processed reads were aligned to the mouse reference genome (mm39) using Bowtie2 (v2.5.1) with default parameters^52^. Reads with mapping quality (MAPQ) ≤ 20, unmapped reads, secondary alignments, or PCR duplicates were filtered out using Sambamba (v1.0.0) ^60^ with the expression: ‘mapping_quality>=20 and ref_name =∼ /^chr[0-9XY]/ and not duplicat’.

Biological replicate concordance was assessed using deepTools ‘plotCorrelation’ (v3.5.1) with Spearman correlation coefficients calculated in 50 kb bins ^61^. Replicates with high correlation coefficients (coef > 0.8) were merged using SAMtools ‘merg’ for downstream analysis.

#### Coverage track generation, and domain calling

Coverage tracks for LMNB1 and H3K9me2 ChIP-seq datasets were generated by extending reads to the estimated fragment length (120 bp) and normalizing to counts per million (CPM). The enrichment of the IP libraries over their corresponding inputs was calculated as a log_₂_(IP/Input) signal in 1 kb bins genome-wide using deepTools bamCoverage and bamCompare. The resulting signal was smoothed with a 10 kb sliding window. Regions present in the ENCODE blacklist for mm10^62^ were lifted over to mm39 using CrossMap (v0.7.3)^63^ and excluded from all analyses. Normalized coverage tracks were visualized using pyGenomeTracks (v3.8)^64^.

Domains of enriched ChIP-seq signal were identified using a 2-state Hidden-Markov Model implemented in HMMDomainCaller^65^ with minor modifications (https://github.com/YiZhang-lab/H2Aub_H3K27me3_preimplantation_dynamics). The HMM was trained on ChIP-seq signal from control samples, using a bin size of 5kb with a 10-bin sliding average. Following HMM domain calling, we applied a post-filtering step in which domains were merged if they were separated by less than 20Kb and discarded if either their length was greater than 10M. LAD calls were compared to previously published datasets using BEDTools (v2.31.1)^66^, BEDOPS (v2.4.41)^67^, eulerR (v7.0-0), and Intervene^68^ to assess concordance.

#### ChIP-seq Signal Enrichment

Mean normalized ChIP-seq signal over domains was computed with deepTools ‘computeMatrix scale-regions’ and visualized using ‘plotProfil’. For genome-wide comparisons, coverage matrices were generated by scaling all domains to their median size. ChIP-seq signal distributions between wildtype and Sun1/2^Pdgfra-KO^ groups were compared using dsCompareCurves (v1.2.0) with paired Wilcoxon signed-rank tests (two-sided, p < 0.05). Statistical robustness was assessed by generating 1000 bootstrap resamples of the paired differences to estimate 95% confidence intervals. Regions passing the Wilcoxon significance cutoff were flagged as differentially occupied. Mean difference curves and confidence intervals were plotted in R (v4.3.2) using ggplot2 (v3.4.4). Domains identified in the wild type group were compared to those from the Sun1/2^Pdgfra-KO^ group using BEDOPS ‘bedmap’ with the ‘--echo --bases --bases-uniq-f’ options to calculate fractional base-pair overlap.

### RNA isolation, cDNA synthesis, and qRT-PCR

Wildtype and Sun1/2^Pdgfra-KO^ fibroblasts were sorted and isolated as described above. Fibroblasts were spun down at 300g for 5 minutes at 4oC. Cells were then lysed with the RNA lysis buffer supplied in the RNeasy Micro Kit (Qiagen, cat# 74004). RNA was isolated according to manufacturer’s protocol using the RNeasy Micro Kit (Qiagen, cat# 74004). RNA concentration was quantified using a NanoDrop spectrophotometer. cDNA was synthesized using the SuperScript VILO IV kit (Thermo Fisher Scientific, cat# 11756050). Quantitative real-time PCR (qRT-PCR) was completed using SYBR reagents (Thermo Fisher Scientific, cat# 4367659) and analyzed using a QuantStudio 7 Pro (Applied Biosystems). All qRT-PCR experiments are analyzed with the delta delta Ct method. Results were further normalized to a random control sample within that experiment. Primer sequences are provided in Supp Table 4.

### Pdgfra+ lung fibroblast isolation and culture

Pdgfra-lineage lung fibroblasts were sorted and isolated as described above. Fibroblasts were grown and maintained in DMEM (Gibco, cat# 11965092), supplemented with 10% FBS (Corning, cat# 35-015-CV) and 1% antibiotic-antimycotic (anti/anti) (Gibco, cat# 15240062). Fibroblasts were grown for one passage and then seeded in 96 well plates (10,000 fibroblasts/well). The next day, fibroblasts were treated with a cocktail to stimulate ECM deposition, consisting of DMEM supplemented with 2% FBS, 10ng/mL Tgfb1 (VWR, cat# 10772-048), and 50ug/mL ascorbic acid (Cayman Chemical, cat# A92902)^69^. Fibroblasts were treated with the cocktail for 3 days, fixed with 2% PFA (Thermo Fisher, cat# J19943.K2) for 20 minutes at room temperature then washed 3x in 1x PBS (5 minutes/wash). Samples were then permeabilized and blocked in blocking buffer (1x PBS supplemented with 5% BSA (Jackson Immuno, cat# 001-000-162) and 0.1% Tween-20 (Sigma, cat# P1379)) for 45 minutes at room temperature. Samples were then incubated with primary antibody (Fibronectin, Santa Cruz, cat# sc-81767) diluted in blocking buffer, overnight at 4oC. The next morning, samples were washed 3x in 1x PBS supplemented with 0.1% Tween-20 (PBS-T), 5 minutes/wash. Samples were then incubated with secondary antibody for 1 hour at room temperature. Samples were washed 3x in PBS-T, 5 minutes/wash. Samples were then imaged using an EVOS microscope (Thermo Fisher) and analyzed within ImageJ.

## Statistical analysis

An unpaired two-tailed t test was used to compare two groups and a one-way ANOVA with Tukey’s adjustment for multiple comparisons was used when comparing multiple groups. Statistical significance was considered when calculated p-values were less than 0.05. Spearman correlation analyses were completed within R. All other statistical analyses were completed using Graphpad Prism 9.

## Acknowledgements

We thank the Flow Cytometry Core Laboratory at the Children’s Hospital of Philadelphia and the Cell and Developmental Biology Microscopy Core at the University of Pennsylvania for their technical assistance. This research was supported by the National Institutes of Health (R00-HL173656 to DLJ, R01-HL164929, R01-HL152194, R01-HL132999, U01-HL148857, R01-HL162683, R01-HL168803 to EEM, R35-HL166663, R01-AG082437, U01-DA052715, R01-GM137425 to RJ).

## Data and code availability

All newly generated genomics data have been deposited into GEO (scRNA-seq: GSE312619, ATAC-seq: GSE312388, and ChIP-seq: GSE312620). Any additional information required to reanalyze the data reported in this paper is available from the lead author upon request.

---

**Figure S1:**
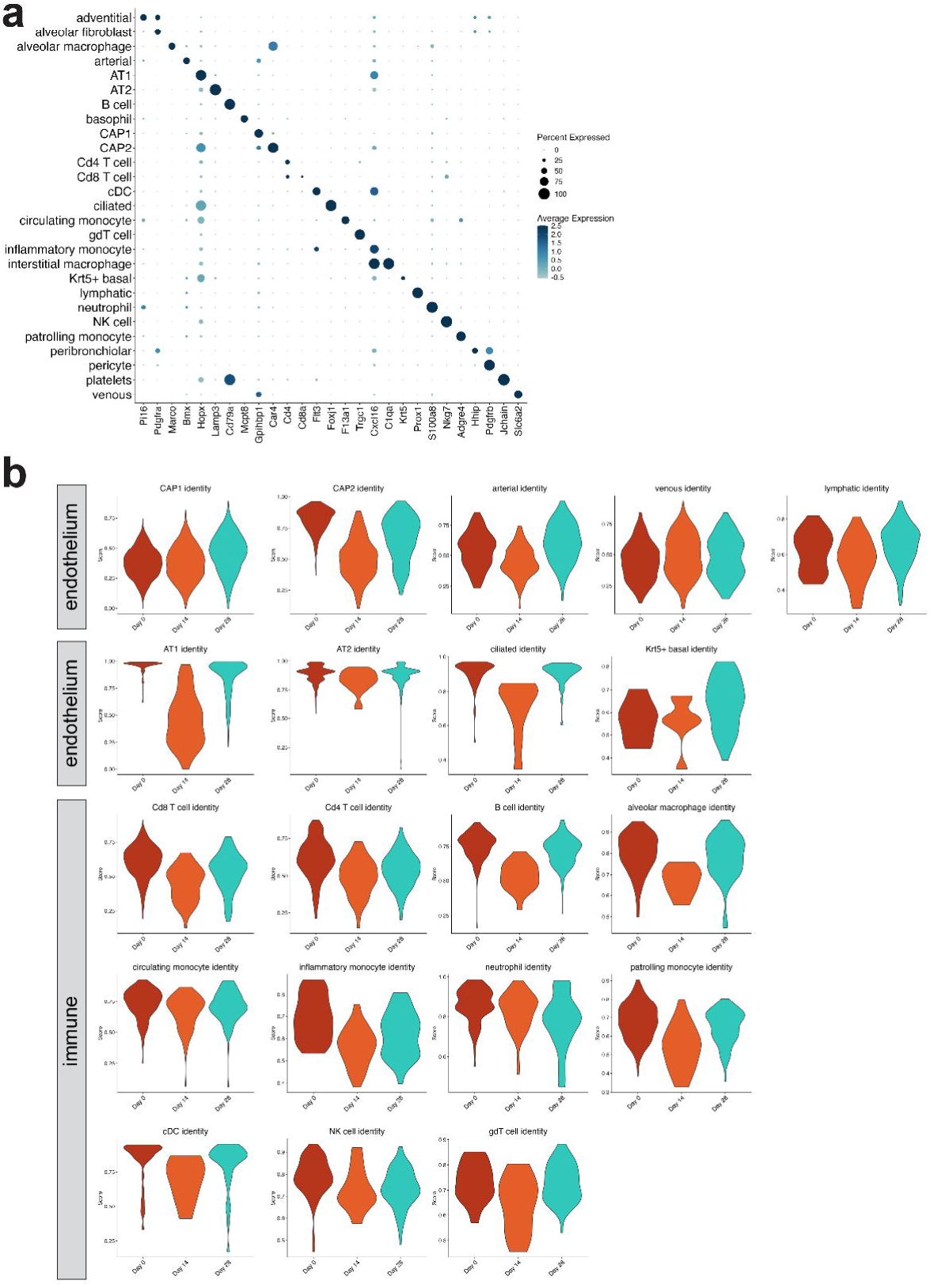
Temporal regulation of injury on lung cell identities. (a) DotPlot showing unique marker expression for each cell type identified in the scRNA-seq dataset. (b) Violin plots of cell type identity scores after bleomycin injury.

**Figure S2:**
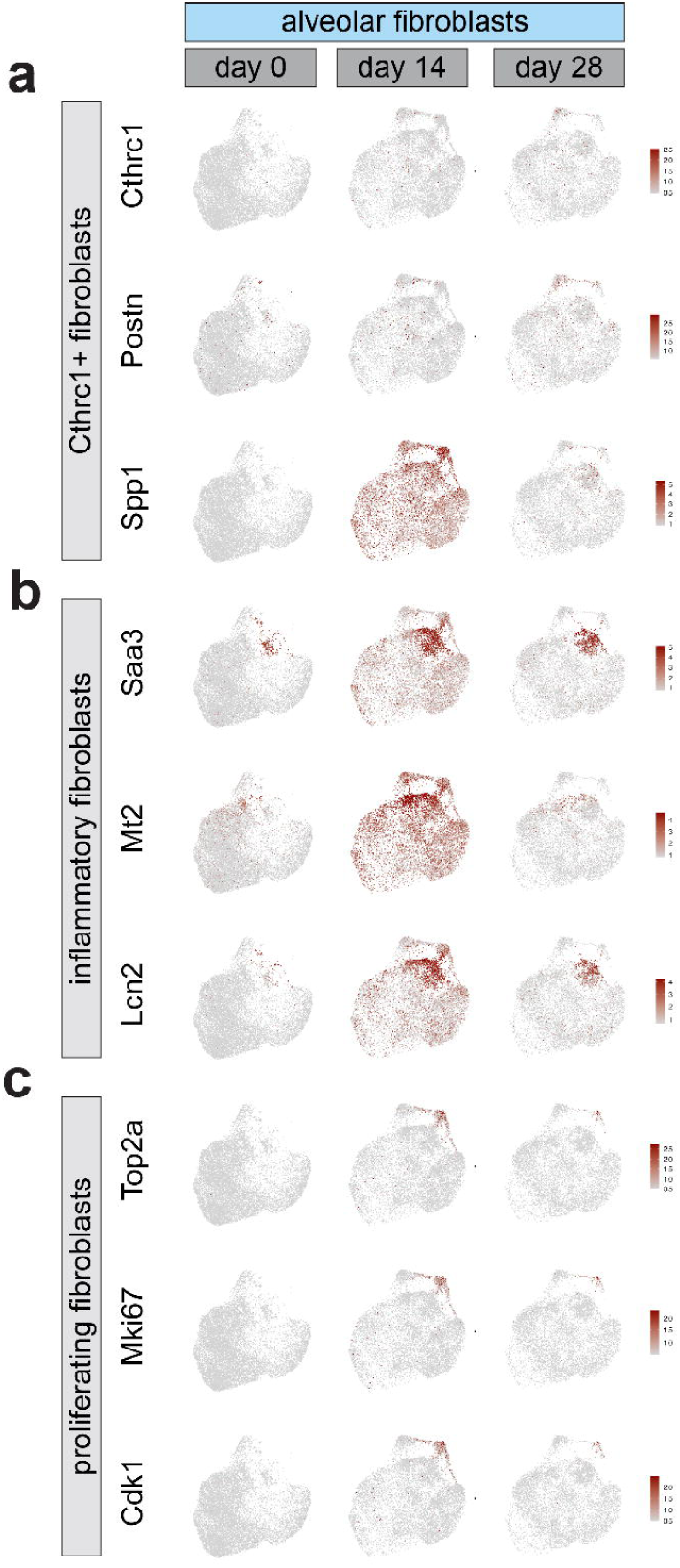
Expression markers of alveolar fibroblast subpopulations after injury. (a) Feature plot demonstrating expression of marker genes of fibrotic Cthrc1+ fibroblasts after injury. (b) Feature plot demonstrating expression of marker genes of inflammatory fibroblasts after injury. (c) Feature plot demonstrating expression of marker genes of proliferating fibroblasts after injury.

**Figure S3:**
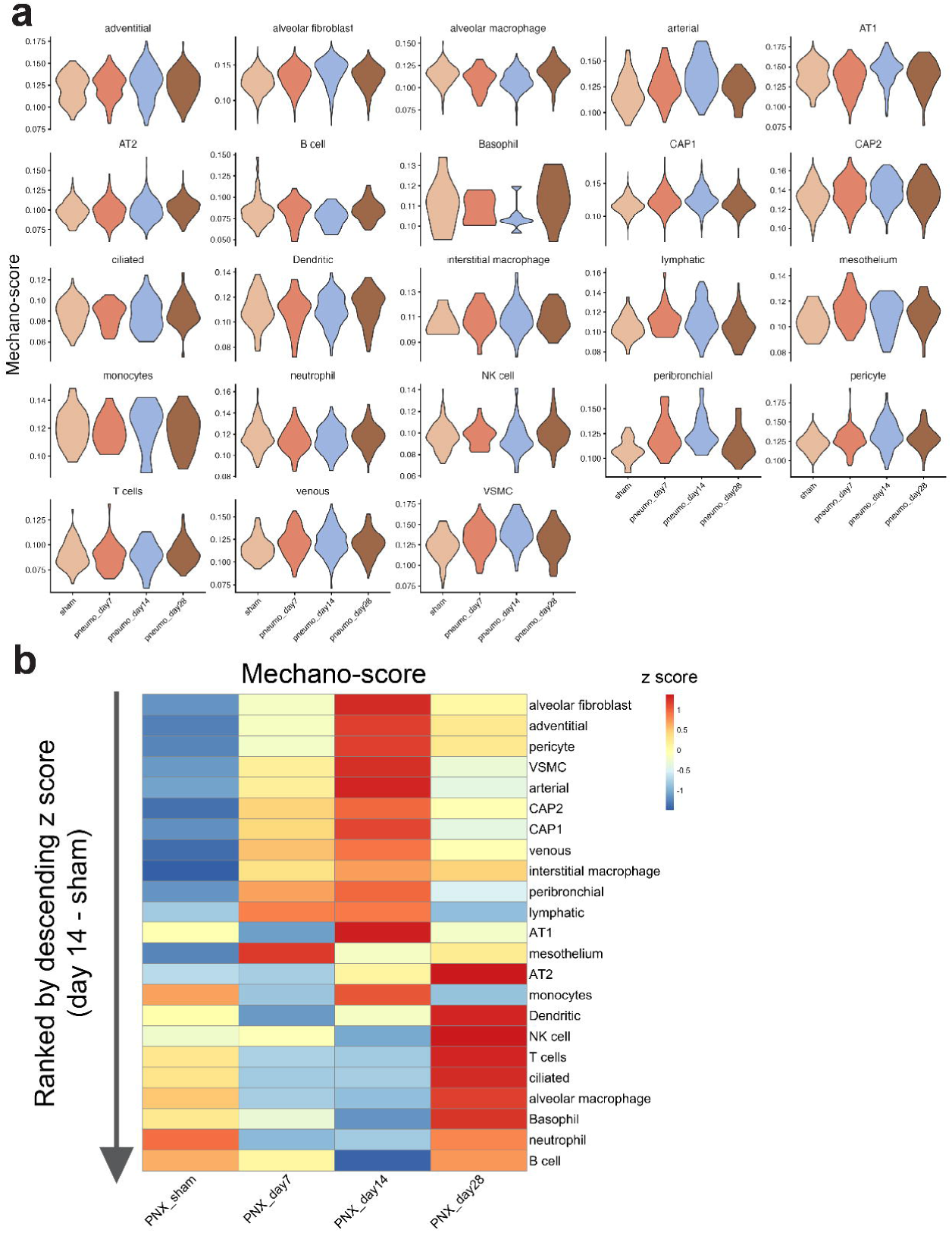
Mechanical regulation of lung cell identity after partial pneumonectomy. (a) Violin plots demonstrating the effect of partial pneumonectomy on lung cell identities over time. (b) heatmap demonstrating mechano-scores for each cell type after partial pneumonectomy. Cells are ranked by descending differential z scores (day 14 – sham).

**Figure S4:**
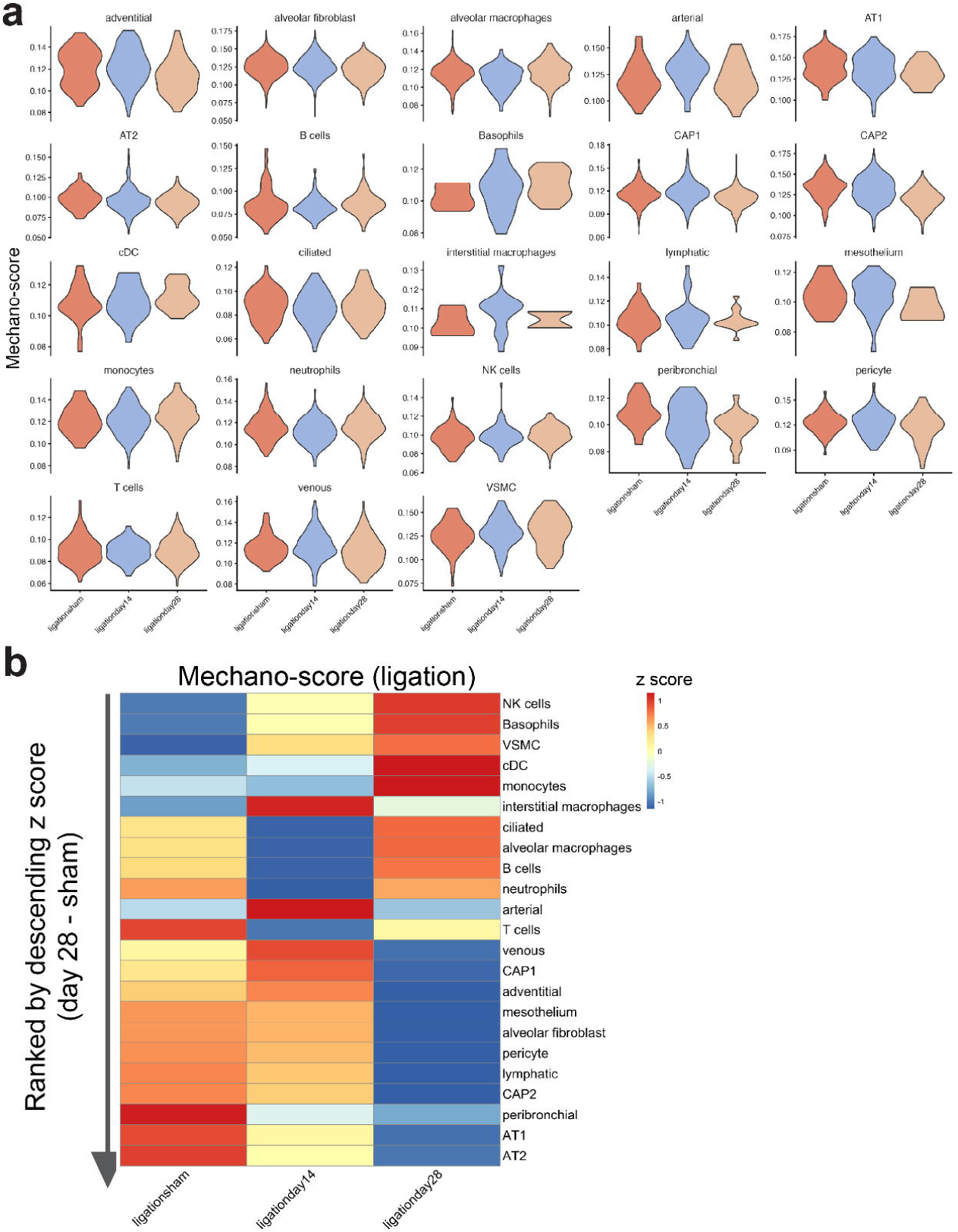
Mechanical regulation of lung cell identity after bronchial ligation. (a) Violin plots demonstrating the effect of bronchial ligation on lung cell identities over time. (b) heatmap demonstrating mechano-scores for each cell type after bronchial ligation. Cells are ranked by descending differential z scores (day 28 – sham).

**Figure S5:**
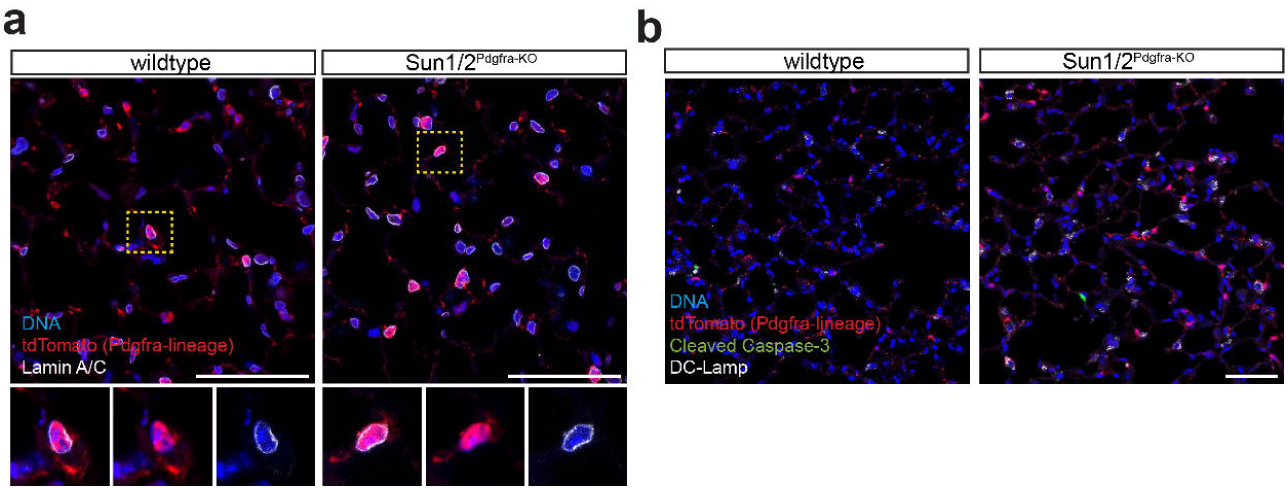
Effect of Sun1 and Sun2 deletion on nuclear morphology. (a) high resolution confocal microscopy image showing Lamin A/C staining in wildtype Pdgfra+ fibroblasts and Sun1/2^Pdgfra-KO^ fibroblasts 4 weeks after Sun1/2 depletion. (b) high resolution confocal microscopy showing Pdgfra+ cells are not apoptotic after Sun1 and Sun2 deletion. Scale bars represent 50 um.

**Figure S6:**
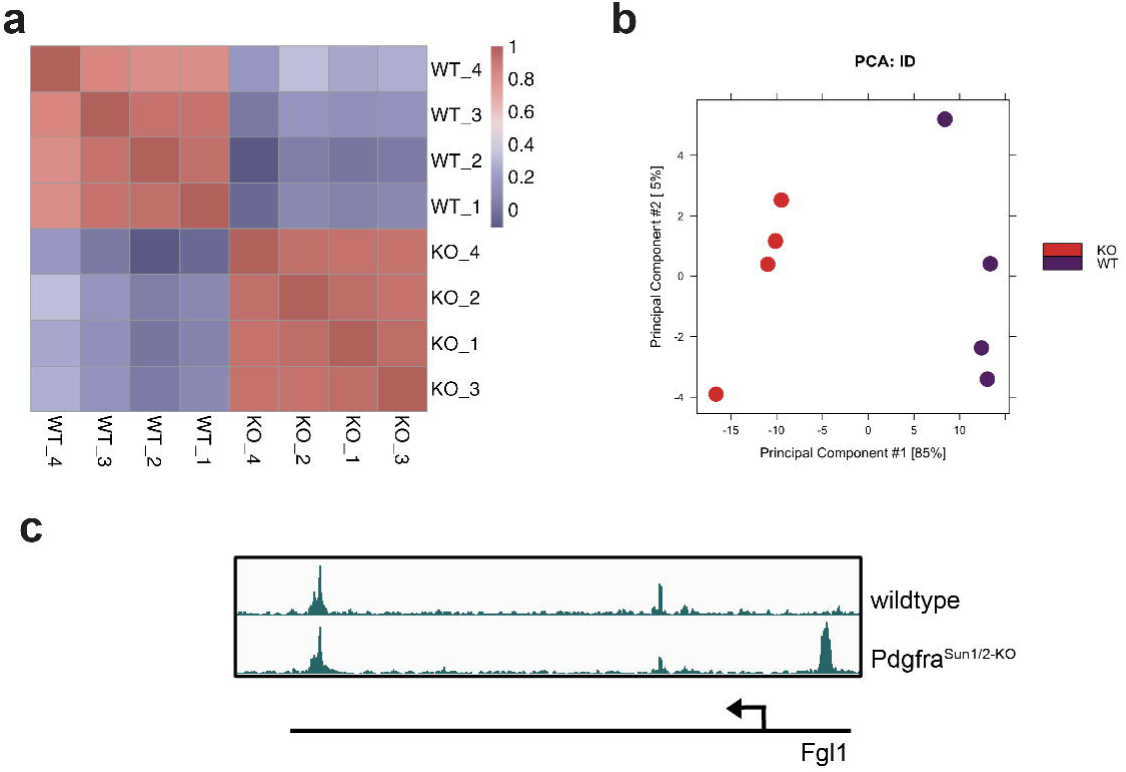
Chromatin accessibility of LINC-depleted lung fibroblasts. (a) Pearson correlation analysis of ATAC-seq data showing that wildtype and Sun1/2^Pdgfra-KO^ fibroblasts are highly correlated within each genotype. (b) Principal component analysis (PCA) demonstrating separation between wildtype and Sun1/2^Pdgfra-KO^ fibroblasts. (c) Representative locus illustrating changes in chromatin accessibility after Sun1/2 depletion in Pdgfra+ fibroblasts.

**Figure S7:**
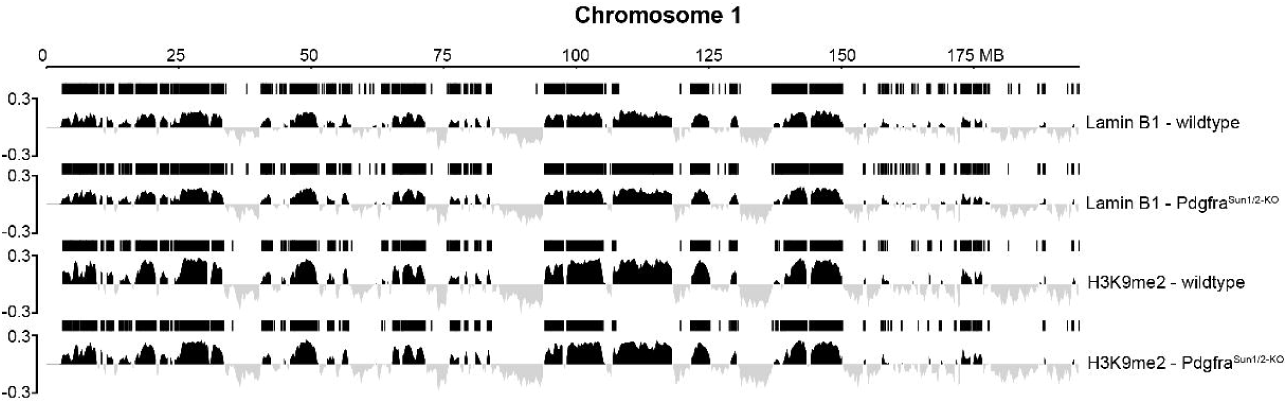
LAD domains of LINC-depleted fibroblasts. Representative genomic track showing LAD domains across chromosome 1 of the mouse genome. LAD domains were assessed by Lamin B1 and H3K9me2 profiles.

**Figure S8:**
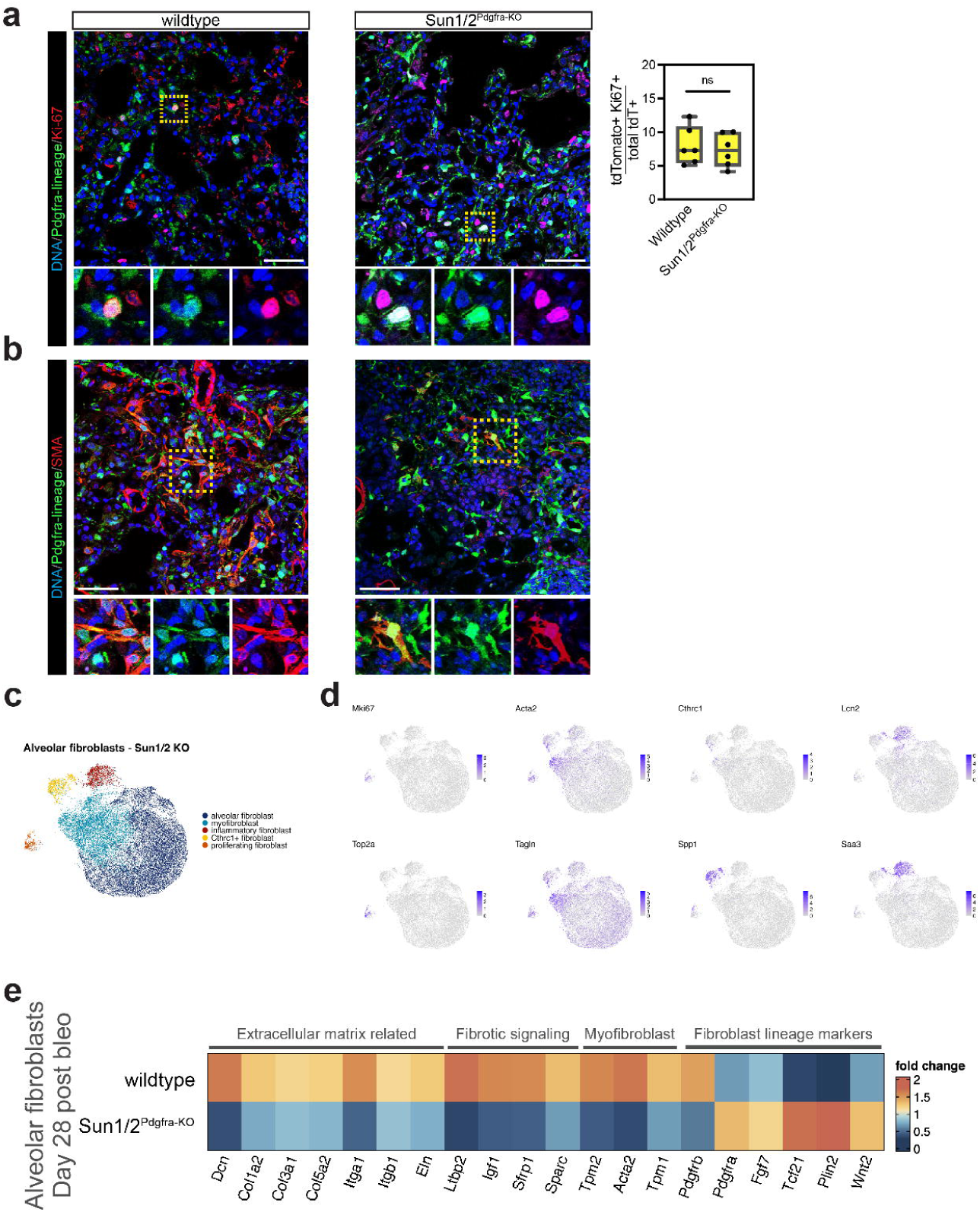
Depletion of the LINC complex in lung fibroblasts does not disrupt their injury response. (a) IHC and quantification showing LINC-depleted Pdgfra+ fibroblasts maintain proliferative potential after bleomycin-induced lung injury. Each dot represents data from a unique biological replicate. Statistical significance evaluated by unpaired t-test. (b) IHC showing LINC-depleted Pdgfra+ fibroblasts maintain capacity to differentiate into myofibroblasts after bleomycin-induced lung injury. All scale bars represent 50 um. (c) UMAP representation of scRNA-seq data of Sun1/2^Pdgfra-KO^ fibroblasts after bleomycin. Data represent merged libraries from sham and days 14 and 28 after bleomycin administration. (d) FeaturePlots showing gene expression of each fibroblast cell state that arises after bleomycin administration (proliferative, myofibroblasts, Cthrc1+ fibroblast, and inflammatory fibroblasts). (e) Heatmap showing expression of differentially expressed genes between wildtype and Sun1/2^Pdgfra-KO^ alveolar fibroblasts at 28 days post bleomycin.

